# Harnessing DNA replication stress to target RBM10 deficiency in lung adenocarcinoma

**DOI:** 10.1101/2023.02.19.529108

**Authors:** Feras E. Machour, Enas Abu-Zhayia, Joyce Kamar, Alma Sophia Barisaac, Itamar Simon, Nabieh Ayoub

## Abstract

The splicing factor RBM10 is frequently mutated in lung adenocarcinoma (LUAD) (9-25%). Most RBM10 cancer mutations are loss-of-function, correlating with increased tumorigenesis and limiting targeted therapy efficacy in EGFR-mutated lung cancer. Notably, therapeutic strategies leveraging RBM10 deficiency remain unexplored. Hence, we conducted RBM10 CRISPR-Cas9 synthetic lethality (SL) screen and identified ∼250 RBM10 SL genes, including WEE1 kinase. WEE1 inhibition sensitized RBM10-deficient LUAD cells *in-vitro* and *in-vivo*. Mechanistically, we identified a splicing-independent role of RBM10 in promoting replication fork progression that underpins RBM10-WEE1 SL. Also, we revealed that RBM10 is associated with active replication forks, which is reliant on PRIM1, an enzyme synthesizing RNA primers for Okazaki fragments. Functionally, we demonstrated that RBM10 serves as an anchor for recruiting HDAC1 and facilitates H4K16 deacetylation to maintain replication fork stability. Collectively, our data revealed a hitherto unrecognized function of RBM10 in fine-tuning DNA replication, and provide therapeutic arsenal for targeting RBM10-deficient tumors.

## Introduction

Lung cancer has the highest mortality rate among all types of cancer worldwide^1,2^. Non-small cell lung cancer (NSCLC) comprises 85% of all lung cancer cases and is subdivided into different subtypes including lung adenocarcinoma (LUAD) (40%)^3,4^. Most LUAD patients are diagnosed at advanced or metastatic stage, when treatment options are limited to surgery, chemotherapy, and few targeted therapies^4^. EGFR and KRAS are the most mutated oncogenes in LUAD amongst Asian (60%) and Caucasian (33%) cohorts, respectively^5,6^. Accordingly, several targeted therapies, including EGFR and KRAS inhibition, are currently used in the clinic for the treatment of advanced stages of LUAD harboring KRAS and EGFR mutations^7,8^. However, effective treatment of LUAD remains elusive due to the genetic diversity of the disease and the development of therapeutic resistance. Thus, there is a critical need for identifying new therapeutic targets for personalized treatment of LUAD. Interestingly, RNA-binding proteins are broadly dysregulated in human cancers including LUAD and play a key role in carcinogenesis and metastatic progression^9–17^. Therefore, alterations in RNA-binding proteins might provide excellent therapeutic targets for treating lung cancer. One very attractive and yet unexplored therapeutic target is the RNA-binding protein, RBM10.

RBM10 is mapped to the X chromosome^18^, and its mutations cause TARP syndrome, an X-linked disorder that leads to pre- and postnatal lethality in affected males^19–25^. RBM10 is a key regulator of alternative splicing that mainly promotes exon skipping of its target genes^26–32^. Also, RBM10 impacts chromatin structure and chromosome segregation independently of its splicing activity^33–36^. Mounting evidence show that RBM10 acts as a tumor suppressor, as its depletion increases cell proliferation and enhances mouse xenograft tumor formation^6,29,32,37–42^. Concordantly, RBM10 overexpression suppresses LUAD tumor growth *in vivo*^6,43^. RBM10 exerts its tumor suppressive activity through various mechanisms^29,40,44^. For example, it was shown that RBM10 inhibits cell proliferation and tumor growth through alternative splicing regulation of NUMB and EIF4H genes^6,32,39^

RBM10 is the most mutated splicing factor in LUAD and is mutated in 9% of all LUAD cases^5^. Strikingly, this percentage rises to 21% in invasive subtypes of LUAD, and 25% in multiple primary LUAD^45,46^. Moreover, RBM10 mutation rate is higher than KRAS and TP53 mutations in early stages of LUAD in patients from Asian origin^47,48^. Most RBM10 mutations in LUAD correspond to loss-of-function and are associated with poor survival^49,50^. Notably, RBM10 mutations mostly co-occur with EGFR and KRAS mutations and were shown to reduce the efficiency of tyrosine kinase inhibitors (TKIs) in LUAD patients harboring EGFR mutations^6,51^. Altogether, these findings highlight the urgency of therapeutic targeting of RBM10 deficiency in LUAD.

Herein, we performed a genome-wide CRISPR-Cas9 synthetic lethality (SL) screen in isogenic LUAD cell line harboring RBM10 cancer mutation and identified ∼250 high-scoring RBM10 SL genes, including WEE1 and Aurora A kinases. We show that pharmacological inhibition of WEE1 selectively sensitizes RBM10-deficient LUAD cells, including patient-derived cells harboring RBM10 cancer mutations, *in vitro* and in mouse xenograft model, and this effect was further exacerbated when combined with Aurora A inhibition. Mechanistically, we identified a splicing-independent role of RBM10 in promoting DNA replication fork progression that underpins RBM10-WEE1 SL. We also show that RBM10 is associated with active replication forks, which is contingent upon PRIM1, an enzyme responsible for synthesizing RNA primers of Okazaki fragments. Functionally, we demonstrated that RBM10 loss disrupts the localization of HDAC1 at replication forks, leading to elevated levels of H4K16ac, which destabilizes the replication fork and induces replication stress. Collectively, our data revealed a hitherto unrecognized function of the RNA-binding protein RBM10 in fine-tuning DNA replication, and identify DNA replication stress as an SL pathway with RBM10 loss. Moreover, we provide a repertoire of RBM10 SL targets that can be harnessed therapeutically to eradicate RBM10-deficient tumors with immediate clinical applicability.

## Results

### Genome-wide CRISPR-Cas9 screen reveals synthetic lethal partners of RBM10 in LUAD cells

Given that RBM10 is one of the most mutated genes in LUAD, we sought to identify genetic vulnerabilities to RBM10 loss. To achieve this, we conducted an unbiased genome-scale CRISPR-Cas9 SL screen in LUAD HCC827 cell line. HCC827 cells harbor a common oncogenic deletion in EGFR (E746-A750), and are a pertinent model to screen for genetic vulnerabilities to RBM10 loss since RBM10 and EGFR mutations often co-occur in LUAD, and RBM10 loss was shown to limit the response to common EGFR-targeted therapy^6,51^. Therefore, we sought to generate isogenic HCC827 cells stably expressing comparable levels of flag-Cas9 endonuclease and differ only in the status of RBM10. Toward this end, we designed a sgRNA targeting RBM10 exon 2 to introduce a frameshift mutation V176Rfs*, that exhibits a complete absence of RBM10 protein, while constitutively expressing Cas9 protein (HCC827-Cas9^RBM10-KO^) (Fig. 1a and Supplementary Fig. 1a,b). Interestingly, the introduced mutation closely resembles a previously characterized LUAD cancer mutation E177* that abrogates RBM10 splicing activity and leads to RBM10 protein loss^50^. Notably, HCC827-Cas9^RBM10-KO^ cells exhibit an increase in exon inclusion of RBM10 splicing targets NUMB and EIF4H, phenocopying the defective splicing phenotype of RBM10-deficient LUAD^6,32^ (Fig. 1b and Supplementary Fig. 1c).

**Figure 1:**
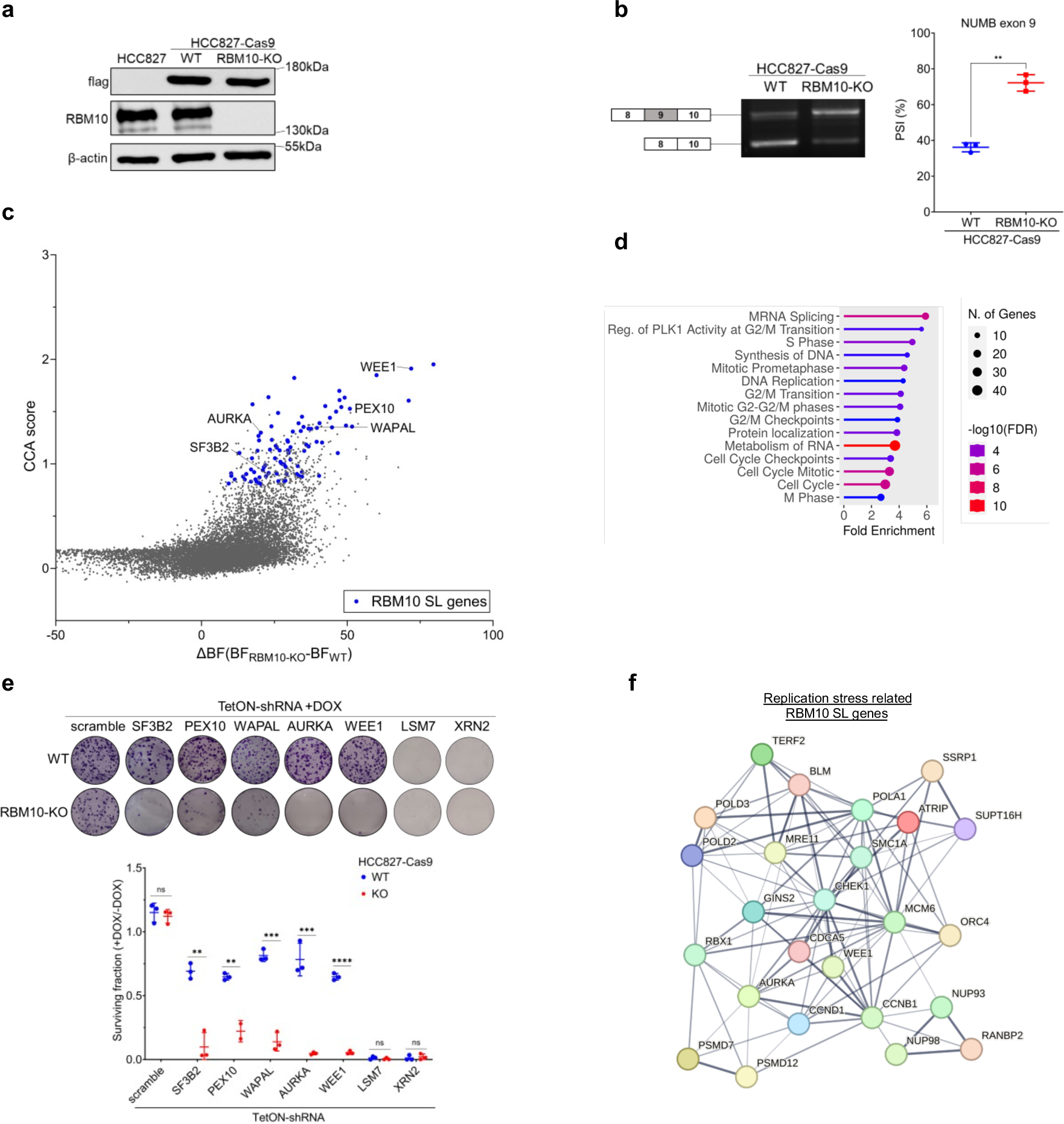
Genome-wide CRISPR-Cas9 screen reveals synthetic lethal partners of RBM10 in LUAD cells. **(a)** Immunoblot analysis for RBM10 and flag-Cas9 protein expression in isogenic RBM10-deficient HCC827 cells constitutively expressing flag-Cas9. β-actin is used as a loading control. The positions of molecular weight markers are indicated to the right. **(b)** RT-PCR analysis of NUMB exon 9 alternative splicing in parental (WT) and HCC827-Cas9^RBM10-KO^ cells. RNA was isolated from the indicated cell lines and analyzed by RT-PCR using primers flanking NUMB exons 8-10. Left: Representative agarose gel image showing amplification of two NUMB variants that differ in exon 9 inclusion. Right: Precent-spliced-in (PSI) quantification of NUMB exon 9 inclusion. Data are presented as mean ± s.d. (n=3). Statistical significance was determined by unpaired t-test*. ***P<0.001*. **(c)** Results of CRISPR-Cas9 SL screen in WT and HCC827-Cas9^RBM10-KO^ cells performed in triplicates. CRISPR Counts Analysis (CCA) score is plotted against the difference in gene essentiality score, expressed as Bayes Factor (BF), between RBM10-KO cells and WT cells. RBM10 SL genes are shown in blue. **(d)** Gene ontology analysis of RBM10 SL genes identified in the CRISPR-Cas9 SL screen. **(e)** Clonogenic survival of WT and HCC827-Cas9^RBM10-KO^ cells transduced with inducible vector expressing either scramble shRNA or shRNA targeting the indicated genes upon the addition of doxycycline (DOX). Top: Representative images of crystal violet staining of the indicated cell lines treated with DOX. Bottom: Quantification of clonogenic survival in DOX-treated cells normalized to untreated cells. Data are presented as mean ± s.d. (n=3). Statistical significance was determined by unpaired t-test. \**\*P<0.01;***P<0.001;****P<0.0001.* **(f)** STRING interaction network of RBM10 SL genes involved in replication stress response and cell cycle. The thickness of the connecting line indicates interaction confidence.

Next, we conducted a genome-scale CRISPR-Cas9 SL screen in the isogenic HCC827-Cas9^WT^ and HCC827-Cas9^RBM10-KO^ using the TKOv1 sgRNA library, which contains 91,320 sgRNAs targeting 17,232 protein-coding genes^52^. Cells were collected at 0 and 15 days following TKOv1 library transduction and subjected to next-generation sequencing to measure sgRNA abundance (Supplementary Fig. 1d). To identify bona fide RBM10 SL genes that are selectively essential in RBM10-KO cells, we analyzed the screen results using three distinct scoring methods: Bayesian Analysis of Gene Essentiality (BAGEL2)^53^, Model-Based Analysis of Genome-wide CRISPR-Cas9 Knockout (MAGeCK)^54^, and CRISPR Count Analysis (CCA)^55^. These methods unveiled 210, 280, and 357 RBM10 SL genes, respectively (Fig. 1c, Supplementary Fig. 1e, and Supplementary Table 1). Of note, a comparative analysis of RBM10 SL genes among the three different assays, revealed 60 RBM10 SL genes that are common to all three analysis methods (Supplementary Fig. 1f). Gene-set enrichment analysis revealed significant enrichment of several pathways including mRNA splicing, cell cycle checkpoints, S-phase, and DNA replication (Fig. 1d).

To functionally validate the screen results, we conditionally knocked-down 5 candidate RBM10 SL genes annotated to various enriched pathways (SF3B1, WAPAL, WEE1, AURKA, PEX10), and 2 common essential (XRN2, LSM7), followed by clonogenic survival assay. While depletion of common essential genes had a similar effect in inhibiting the proliferation of both parental and RBM10-KO cells, depletion of candidate RBM10 SL genes selectively inhibited the proliferation of HCC827-Cas9^RBM10-KO^ cells, confirming successful identification of bona-fide RBM10 SL genes (Fig. 1e and Supplementary Fig. 1g).

Interestingly, several RBM10 SL genes, such as WAPAL, WEE1, CHEK1, ATRIP, and MRE11, were identified using the three aforementioned scoring methods. These genes have known functions in DNA replication stress response and replication fork protection, suggesting DNA replication stress as a synthetic lethal pathway with RBM10 deficiency in LUAD (Fig. 1f). In support of this, RBM10 was previously identified as a candidate replication stress response gene by a genome-wide siRNA screen^56^, and a phosphorylation target of ATR^57^. Additionally, RBM10 was shown to interact with several proteins involved in DNA replication and repair in fission yeast^36^. Fittingly, analysis of “The Cancer Genome Atlas” (TCGA^58^) mutation data from LUAD patients revealed that RBM10 deficiency is associated with increased tumor mutational burden and replication stress gene signature (Supplementary Fig. 1h,i). Collectively, these observations suggest a role of RBM10 in DNA replication and genomic stability.

### RBM10 promotes DNA replication fork progression and replication stress response

Herein, we sought to determine the impact of RBM10 loss on replication fork progression. DNA fiber analysis showed that HCC827-Cas9^RBM10-KO^ cells display a significant replication fork slowdown, suggesting that RBM10 is required for proper fork progression (Fig. 2a). Similar results were also observed in a second RBM10-KO clone (KO2) isolated from HCC827-Cas9 cells, arguing against a clonal effect (Supplementary Fig. 2a). To confirm that the effect of RBM10 loss on replication fork progression is not cell-line specific, and since TP53 mutations often co-occur with RBM10 mutations in LUAD^58^, we tested the effect of RBM10 knockout on replication fork progression in p53-deficient LUAD cells NCI-H1299 (H1299^RBM10-KO^) (Supplementary Fig. 2b,c). Results show that RBM10 loss in H1299 cells also led to significant reduction in replication fork progression, suggesting that the effect of RBM10 on DNA replication is independent of common co-occurring mutations such as EGFR and TP53 (Fig 2b). Immunostaining analysis revealed that replication fork slowdown upon RBM10 loss is accompanied by an increase in the levels of pRPA32-S33 and γH2AX, primarily observed in S phase cells marked by EdU staining (Fig. 2c-e). Similarly, western blot analysis demonstrated a significant increase in pRPA32-S33, pCHK1, and γH2AX in RBM10 knockout (KO) cells synchronized at the S phase using a double-thymidine block (Supplementary Fig. 2d). Next, we investigated the effect of RBM10 loss on the recovery from hydroxyurea (HU) induced replication stress. Results showed that RBM10-deficient cells display a significant increase in replication stress and DNA damage markers after release from HU, with no significant effect on replication fork restart (Fig. 2f and Supplementary Fig. 2e). Corollary to this, RBM10-deficient cells are markedly more sensitive than RBM10-proficient cells to increasing HU concentrations (Fig. 2g). Collectively, our results demonstrate that RBM10 fosters efficient DNA replication fork progression and fork recovery during replication stress.

**Figure 2:**
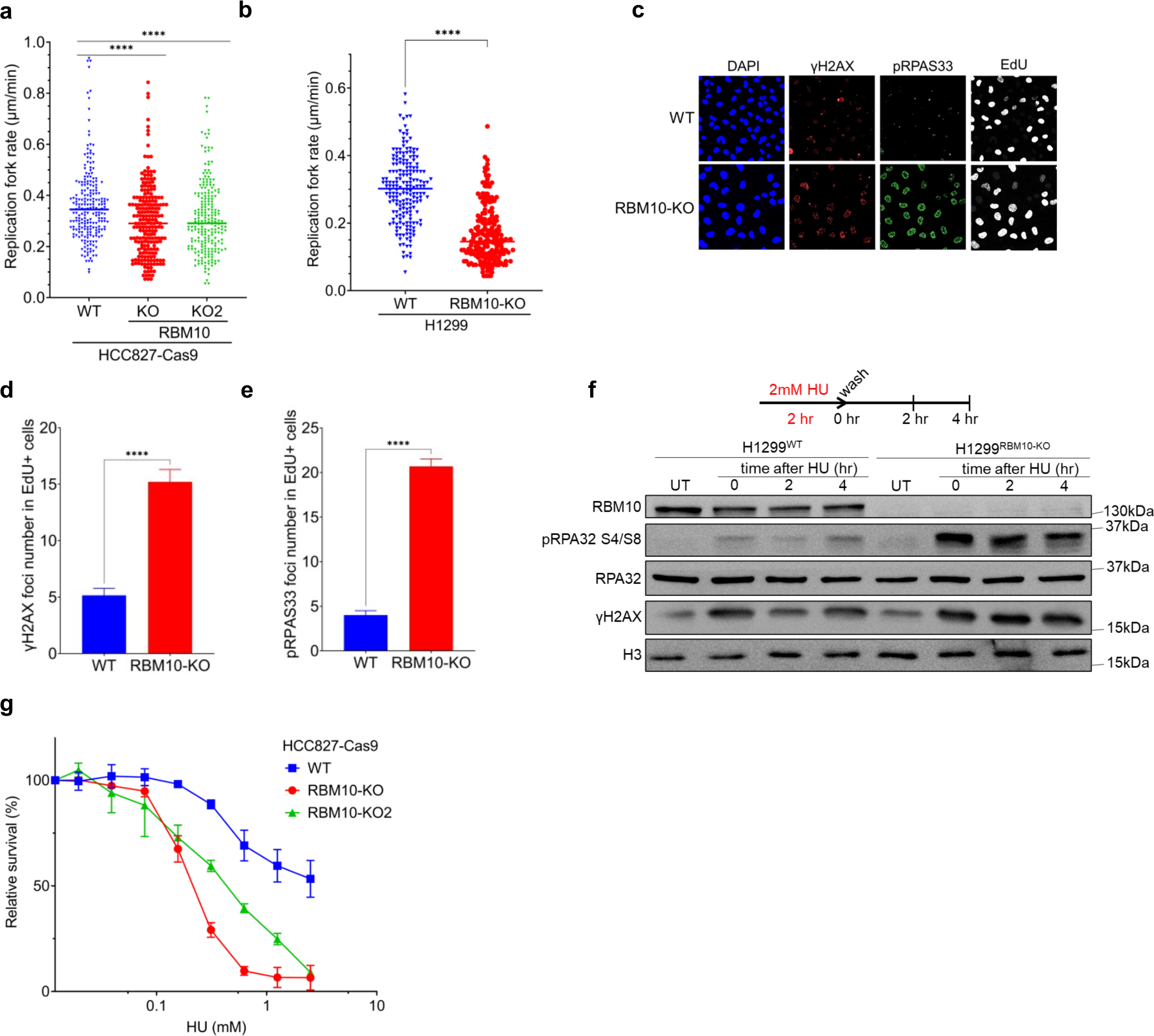
RBM10 promotes DNA replication fork progression and replication stress response. **(a,b)** Replication fork speed measurement using DNA combing assay in WT and RRBM10-KO HCC827 (a) or H1299 (b) cells. Horizontal bars represent mean value of replication fork speed (n>200). Statistical significance was determined by Mann-Whitney test. *\*\*\*\*P<0.0001*. **(c)** Representative immunofluorescence microscopy images of pRPA32-S33 and γH2AX foci in H1299^WT^ and H1299^RBM10-KO^ cells. EdU is used to mark S-phase cells. DAPI is used to stain nuclei. Scale bar, 20µm. **(d,e)** Quantification of γH2AX (d) and pRPA32-S33 (e) foci in S-phase (EdU-positive) H1299^WT^ and H1299^RBM10-KO^ cells. Data are presented as mean foci number per nucleus ± SEM (n>200 cells) and representative of 3 independent experiments. Statistical significance was determined by Mann-Whitney test. *\*\*\*\*P<0.0001*. **(f)** H1299^WT^ and H1299^RBM10-KO^ cells were treated with 2mM hydroxyurea (HU) for 2 hr and subjected to immunoblot analysis at the indicated times after release from HU. Histone H3 is used as a loading control. UT=untreated. The positions of molecular weight markers are indicated to the right. **(g)** Short-term cell viability assay in parental (WT) and RBM10-KO HCC827-Cas9 cells treated with increasing concentrations of HU. Data are presented as mean ± s.d. (n=3).

### RBM10 is associated with active DNA replication fork in a PRIM1-dependent manner

To gain molecular insights into the involvement of RBM10 in replication fork progression, we first determined its sub-cellular localization. Biochemical fractionation revealed that RBM10 is enriched at the chromatin-bound fraction (Supplementary Fig. 3a), supporting the notion that RBM10 has a direct role in regulating replication fork progression. To further substantiate this, cells expressing RBM10 fused to EGFP were subjected to GFP-trap followed by mass spectrometry (MS) analysis to map RBM10 interactome. Results revealed 410 RBM10-interacting proteins annotated to several pathways, most notably the spliceosome, consistent with RBM10 function in alternative splicing, as well as DNA replication. (Fig. 3a and Supplementary Table 2). Intriguingly, we revealed that RBM10 interacts with replication fork components and proteins involved in replication stress response (Supplementary Fig. 3b). Next, we validated the authenticity of RBM10 interactome and showed that RBM10 interacts with MCM5, PCNA, HDAC1, PARP1, and RAD51 (Fig. 3b).

**Figure 3:**
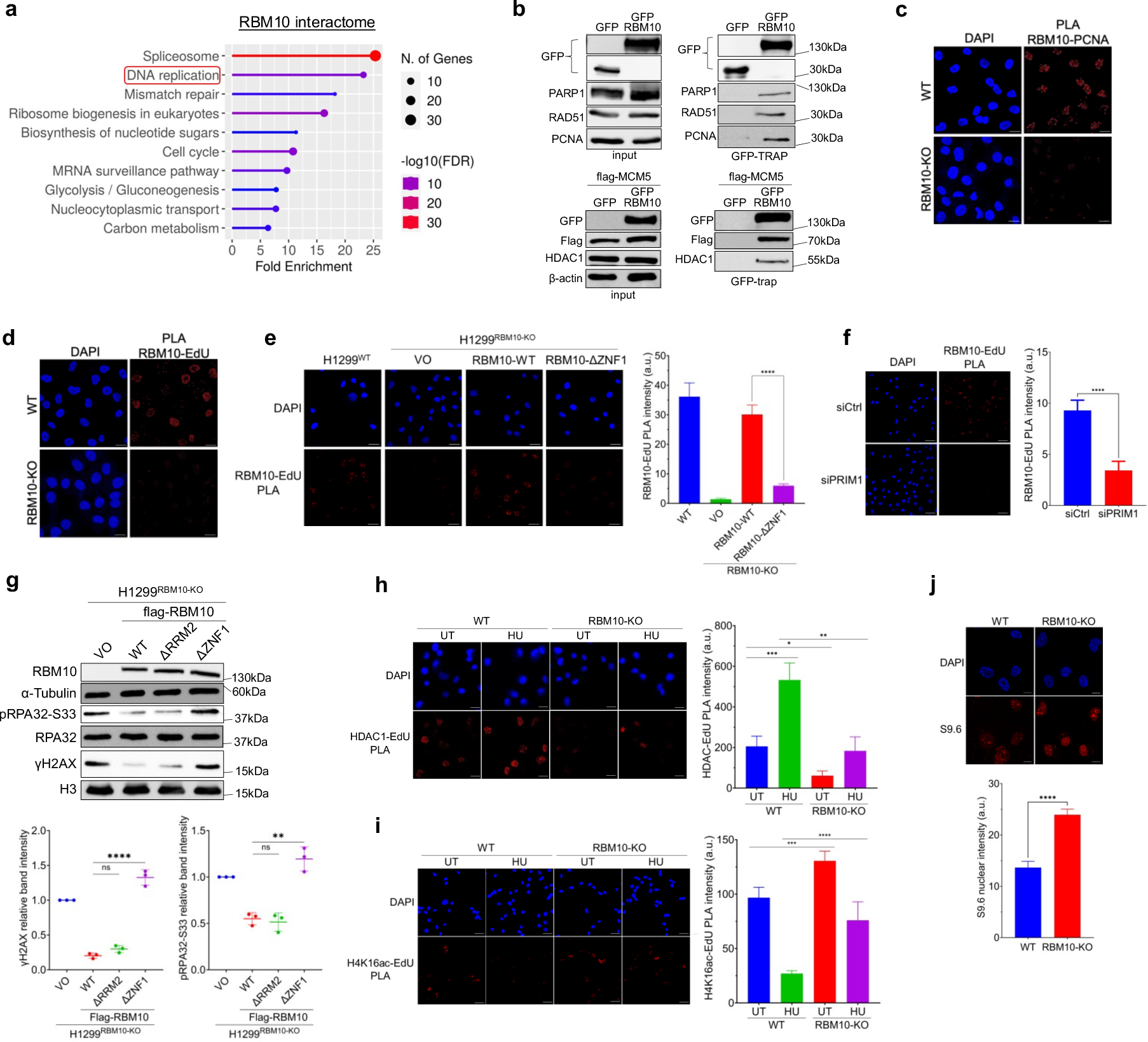
PRIM1-dependent RBM10 association with DNA replication forks promotes HDAC1 recruitment to limit replication stress. **(a)** Gene ontology analysis of RBM10 interactome in unchallenged cells as identified by GFP-trap followed by mass spectrometry analysis. **(b)** HEK293T cells expressing EGFP-RBM10 or EGFP only were subjected to GFP-trap followed by immunoblot analysis using the indicated antibodies. Bottom: GFP-trap was performed on HEK293T cells additionally expressing Flag-MCM5. The positions of molecular weight markers are indicated to the right. **(c,d)** Representative immunofluorescence microscopy images of RBM10:PCNA (c) and RBM10:EdU-biotin (d) proximity ligation assay (PLA) foci in H1299^WT^ and H1299^RBM10-KO^ cells. Scale bar, 20µm. **(e)** Left: Representative immunofluorescence microscopy images of RBM10:EdU-biotin PLA in H1299^RBM10-KO^ expressing Flag-RBM10^WT^, Flag-RBM10^ΔZNF1^, or vector only (VO). DAPI is used to stain nuclei. Scale bar, 20µm. Right: Quantification of RBM10-EdU-biotin PLA intensity per nucleus. Data are presented as mean PLA intensity per nucleus ± SEM (n>300 cells). Statistical significance was determined by Mann-Whitney test. *\*\*\*\*P<0.0001*. **(f)** As in (e), except H1299^WT^ were transfected with siRNA against PRIM1 (siPRIM1) or control siRNA (siCtrl) 72 hr prior to PLA assay. Scale bar, 50µm. **(g)** Immunoblot analysis of H1299^RBM10-KO^ cells expressing Flag-RBM10^WT^, Flag-RBM10^ΔRRM2^, Flag-RBM10^ΔZNF1^, or vector only (VO). Top: Representative immunoblot image using the indicated antibodies. α-Tubulin is used as a loading control. Bottom: Quantification of γH2AX band intensity (relative to H3) and pRPA32-S33 band intensity (relative to RPA32). Data are presented as mean ± s.d. (n=3). Statistical significance was determined by unpaired t-test. \**\*P<0.01;****P<0.0001;ns, not significant*. **(h,i)** Left: Representative immunofluorescence microscopy images of HDAC1:EdU-biotin (h) and H4K16ac:EdU-biotin (i) PLA in H1299^WT^ and H1299^RBM10-KO^ cells. Cells were labelled with EdU for 20 min and were either left untreated (UT), treated with 1 mM HU for 1 h (HU). DAPI is used to stain nuclei. Scale bar, 20µm. Right: Quantification of PLA intensity per nucleus. Data are presented as mean PLA intensity per nucleus ± SEM (n>50). Statistical significance was determined by Mann-Whitney test. *\*P<0.05;**P<0.01;***P<0.001*. **(j)** Top: Representative immunofluorescence microscopy images of R-loops detected by S9.6 antibody in H1299^WT^ and H1299^RBM10-KO^ cells. Cells overexpressing RNaseH1 are used as a negative control. DAPI is used to stain nuclei. Scale bar, 10µm. Bottom: Quantification of S9.6 signal in H1299^WT^ and H1299^RBM10-KO^ cells. Data are presented as mean intensity per nucleus ± SEM (n>50 cells) and representative of 3 independent experiments. Statistical significance was determined by Mann-Whitney test. *\*\*\*\*P<0.0001*.

To further substantiate RBM10 interaction with replication forks, first we confirmed the interaction between endogenous RBM10 and PCNA in LUAD cells (Supplementary Fig. 3c). Second, we tested whether RBM10 is associated with active replication forks in cells using proximity ligation assay (PLA). Results showed that RBM10 is in close proximity to PCNA (Fig. 3c). Moreover, PLA combined with EdU click chemistry revealed that RBM10 is associated with EdU-labelled nascent DNA, marking sites of active DNA replication (Fig. 3d). Collectively, our data provide firm evidence that RBM10 interacts with active DNA replication forks under native conditions.

Next, we aimed to gain molecular insights into the mechanism governing the association of RBM10 with DNA replication forks. Given that RBM10 is a well-established RNA-binding protein, we hypothesized that its binding to RNA molecules facilitates its association with chromatin. To explore this hypothesis, we treated cells with RNase A followed by chromatin fractionation. Our results revealed that RNase A treatment profoundly disrupts RBM10 association with chromatin (Supplementary Fig. 3d). Consistent with this observation, an RBM10 deletion mutant lacking the first ZnF domain (RBM10^ΔZNF1^), known to bind RNA molecules^59,60^, lost its association with active DNA replication forks, as determined by PLA (Fig. 3e). Remarkably, a recent work has shown that PRIM1, an enzyme involved in synthesizing RNA primers for Okazaki fragments, facilitates the recruitment of 53BP1 to DNA replication forks^61^. Hence, we investigated whether a similar scenario exists with RBM10. Our results reveal that siRNA knockdown of PRIM1 pronouncedly diminishes the association of RBM10 with active DNA replication forks similar to 53BP1 (Fig. 3f and Supplementary Fig. 4a,b). Collectively, these findings robustly support the notion that Okazaki RNA primers play a pivotal role in mediating the recruitment of RBM10 to replication forks.

### The role of RBM10 in replication stress response is distinct from its splicing activity

Since RBM10 modulates the splicing of hundreds of genes, we sought to test whether the splicing activity of RBM10 underlies its role in the DNA replication stress response. To address this, we reintroduced RBM10-deficient cells with a vector expressing either flag-RBM10-WT or RBM10^ΔZNF1^ or RBM10 mutant that lack RRM2 domain (RBM10^ΔRRM2^), known to modulate RBM10 splicing activity^60^. Subsequently, we assessed the ability of these RBM10 mutants to restore both splicing efficiency and the levels of replication stress markers to normal levels. Our results demonstrate that while RBM10^ΔZNF1^ successfully restores the splicing efficiency of RBM10 target genes, NUMB and EIF4H, it still exhibits increased levels of replication stress markers compared to cells expressing RBM10-WT (Fig. 3g and Supplementary Fig. 4c,d). Conversely, RBM10-deficient cells expressing RBM10^ΔRRM2^ mutant show defects in the splicing of RBM10 target genes but display normal levels of replication stress markers compared to control cells. These findings collectively support the notion that the role of RBM10 in DNA replication is independent of its splicing activity. To further validate this observation, we conducted transcriptome analysis of RBM10 in isogenic HCC827 and NCI-H1299 cells. Results revealed no discernible changes in the expression of core DNA replication and repair genes (Supplementary Fig. 4e and Supplementary Table 2). Our findings align with previously reported transcriptome analyses of RBM10 in various cell lines, demonstrating that RBM10 does not regulate the expression or splicing of core DNA replication and repair genes^6,32^.

### RBM10 targets HDAC1 to active replication fork to fine-tune histone acetylation

To elucidate the mechanistic role of RBM10 in DNA replication progression and replication stress response, we focused on RBM10-HDAC1 interaction that has been identified in the RBM10 interactome and validated through immunoprecipitation (Fig. 3b and Supplementary Fig. 3b). Also, fission yeast Rbm10 was previously shown to interact with Clr6, a homolog of human HDAC1/2^36^. In addition, previous reports have shown that HDAC1 is localized at active DNA replication forks, where it deacetylates histones to prevent replication fork collapse^62–65^. Prompted by this, we sought to investigate whether RBM10 is essential for recruiting HDAC1 to DNA replication forks. Our results demonstrated a pronounced disruption in the association of HDAC1 with DNA replication in RBM10-deficient cells under normal and stress conditions (Fig. 3h). Consequently, the levels of H4K16 acetylation were increased in RBM10-deficient cells compared to control cells (Fig. 3i). In summary, these data suggest that RBM10 serves as an anchor for recruiting HDAC1 to active replication forks to facilitate the deacetylation of H4K16, a modification recently recognized as crucial for replication fork stability^65,66^. Since a recent work demonstrated that histone deacetylation counteracts the levels of R-loops at active replication forks to mitigate transcription-induced replication stress^66^, we postulated that the increased levels of H4K16ac in RBM10-deficient cells is also accompanied by increased R-loops levels. Indeed, we observed that RBM10-deficient cells exhibit elevated levels of R-loops in S-phase cells, indicating that replication stress observed upon RBM10 loss is mediated, at least in part, by transcription-replication conflicts and R-loop accumulation (Fig. 3j and Supplementary Fig. 4f,g).

Additionally, since RBM10 interacts with RAD51 protein (Figure 3b) which has a central role in homology repair (HR) of double-strand breaks (DSB) and its depletion impairs HR repair leading to replication stress^67,68^, we sought to investigate the functional relevance of their interaction^67,68^. We aimed to determine whether the replication stress in RBM10-deficient cells is due to defective HR repair. Toward this, we assessed the efficacy of HR repair of DSBs using the Cas9-mClover-LMNA assay^69^. Results indicate that RBM10 deficiency has no discernible effect on the percentage of mClover-positive cells (representing cells that repaired DSBs through HR) compared to control cells (Supplementary Fig. 4h). This observation suggests that the replication stress observed in RBM10-deficient cells is not attributed to defective HR repair. This data raises an interesting possibility that the RBM10-RAD51 interaction might be required for replication fork reversal rather than HR repair.

### WEE1 kinase inhibition selectively sensitizes RBM10-deficient LUAD cells

Herein, we sought to exploit the role of RBM10 in replication stress to selectively target RBM10-deficient LUAD cells. Towards this, we focused on WEE1 kinase, since it is a high scoring RBM10 SL gene that was identified in the three scoring methods (BAGEL, MAGeCK and CCA) (Fig. 1b and Supplementary Fig. 1e,f), and is involved in several RBM10 SL pathways, such as S-phase and G2/M cell cycle checkpoints, and was shown to protect the stability of stalled replication forks^70–72^. Moreover, WEE1 has a small molecule inhibitor, MK1775 (AZD1775), which is currently in several phase II clinical trials for treatment of advanced solid tumors (ClinicalTrials.gov), highlighting the clinical potential of targeting RBM10-deficient LUAD^73–75^. To test whether the observed RBM10-WEE1 synthetic lethality (Fig. 1e) is dependent on WEE1 kinase activity, we pharmacologically inhibited WEE1 using MK1775 in parental HCC827-Cas9^WT^ and two HCC827-Cas9^RBM10-KO^ cell lines. Short and long-term survival assays revealed that HCC827-Cas9^RBM10-KO^ cells exhibit pronounced sensitivity to MK1775 treatment compared to parental cells (Fig. 4a,b, Supplementary Fig. 5a). The hypersensitivity of RBM10-deficient cells to MK1775 was also observed in RBM10-KO (KO3) cells derived from naïve HCC827 cells that do not stably express Cas9, suggesting that RBM10-WEE1 synthetic lethality is not due to a clonal effect or cytotoxicity caused by stable expression of Cas9 endonuclease (Supplementary Fig. 5b,c). In addition, complementing RBM10-deficient cells with a vector expressing flag-RBM10-WT suppressed the sensitivity to MK1775 compared to cells expressing empty vector (Fig. 4c).

**Figure 4:**
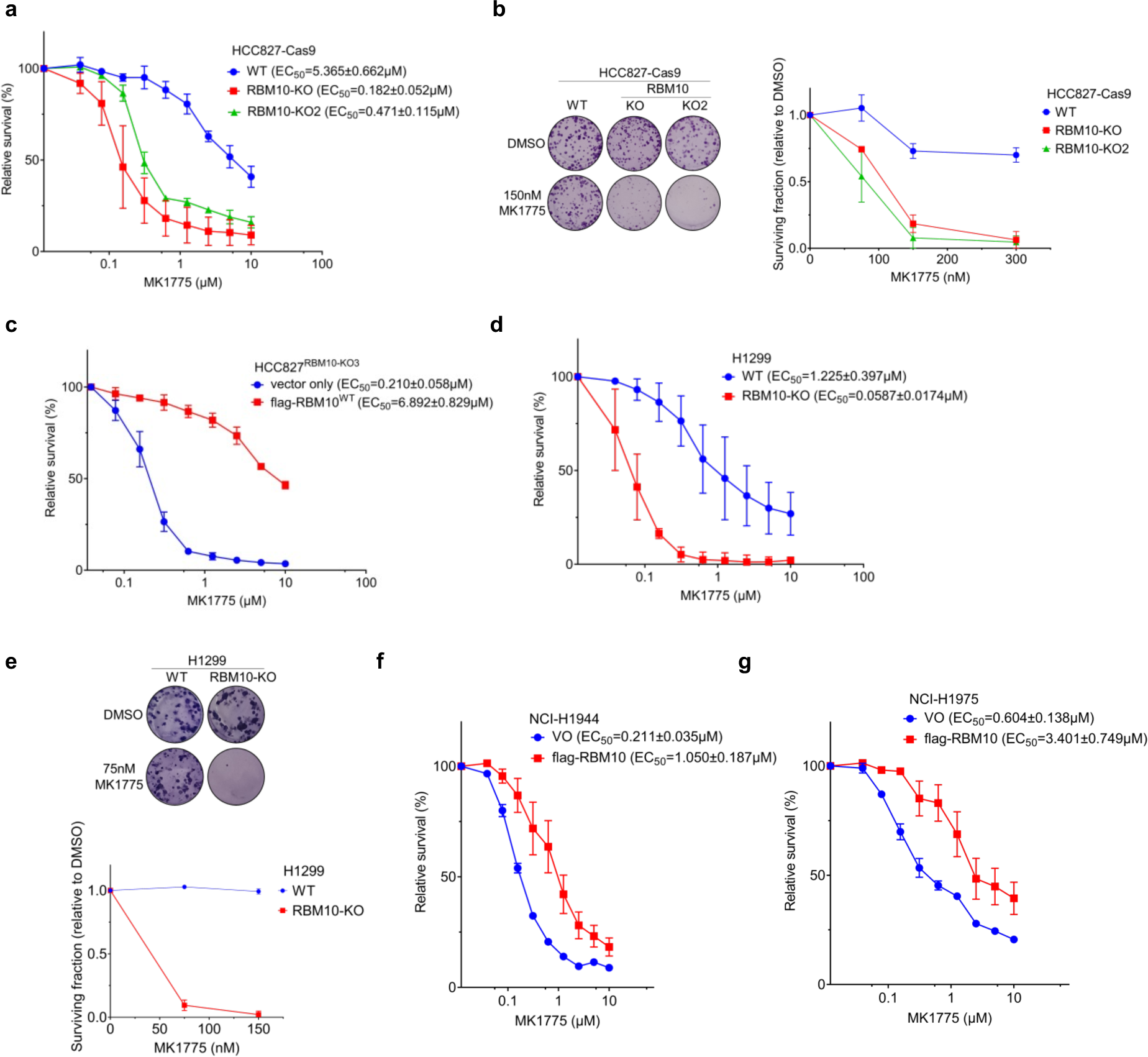
WEE1 kinase inhibition selectively sensitizes RBM10-deficient LUAD cells. **(a)** Short-term cell viability assay and EC_50_ determination in WT and two RBM10-KO clones (KO and KO2) of HCC827-Cas9 cells treated with increasing concentrations of MK1775. Data are presented as mean ± s.d. (n=3). **(b)** Clonogenic survival of WT and two RBM10-KO clones treated with the indicated concentrations of MK1775. Left, representative images of plates stained with crystal violet. Right, quantification of clonogenic survival. Data are presented as mean ± s.d. (n=3). **(c)** As in (a), except of using HCC827^RBM10-KO3^ cells expressing flag-RBM10-WT or vector only. (**d**) As in (a), except of using H1299^WT^ and H1299^RBM10-KO^ cells. **(e)** Clonogenic survival of H1299^WT^ and H1299^RBM10-KO^ cells treated with the indicated concentrations of MK1775. Top: Representative images of plates stained with crystal violet. Bottom: Quantification of clonogenic survival. Data are presented as mean ± s.d. (n=3). **(f,g)** As in (a), except of using NCI-H1944 (f) or NCI-H1975 (g) cells expressing either flag-RBM10-WT or vector only.

Similar to p53-proficient HCC827, our results showed that H1299^RBM10-KO^ cells are hypersensitive to MK1775 compared to control cells (Fig. 4d,e). To increase the therapeutic potential of our observations, we tested RBM10-WEE1 synthetic lethality in two RBM10-deficient patient-derived cell lines, NCI-H1944 (containing KRAS^G13D^ mutation) and NCI-H1975 (containing EGFR^L858R/T790M^ mutation) harboring p.A683Rfs*10 and p.G905Afs*7 truncating RBM10 mutations, respectively. Towards this, NCI-H1944 and NCI-H1975 cells were transduced with flag-RBM10 expression vectors (Supplementary Fig. 5d,e) and treated with MK1775 followed by cell viability assays. Results showed that RBM10 expression rescues the sensitivity of RBM10-deficient cell lines to MK1775, suggesting that RBM10 loss occurring during LUAD tumorigenesis potentiates the sensitivity to WEE1 inhibition (Fig. 4f,g). Altogether, our results recapitulate RBM10-WEE1 synthetic lethality observed in our CRISPR-Cas9 SL screen and demonstrate that WEE1 inhibition selectively sensitizes RBM10-deficient LUAD tumor cells from different origins and independently of common co-occurring mutations such as EGFR, KRAS and TP53, highlighting the clinical relevance of targeting RBM10 loss in LUAD using MK1775.

### DNA damage and premature mitotic entry underpin RBM10-WEE1 synthetic lethality

We sought to decipher the molecular basis underlying the hypersensitivity of RBM10-deficient cells to WEE1 inhibition. We showed that H1299^RBM10-KO^ cells treated with increasing concentrations of MK1775 exhibit elevated levels of phosphorylated histone H2A.X (γH2AX), suggesting that DNA damage accumulation upon WEE1 inhibition is exacerbated in RBM10-deficient cells (Fig. 5a). Likewise, higher levels of stalled forks and replication stress markers including phosphorylated RPA32 on positions 4 and 8 (pRPA32-S4/S8), RPA32 on position 33 (pRPA32-S33), and CHK1 on position 345 (pCHK1-S345) were observed in MK1775-treated H1299^RBM10-KO^ cells compared to control cells (Fig. 5b and Supplementary Fig 6a,b). Since WEE1 inhibition in RBM10-deficient cells leads to excessive replication stress, we investigated the effect of MK1775 treatment on replication fork progression. Results showed that WEE1 inhibition exacerbates the decrease in replication fork rate in RBM10-deficient cells compared to parental cells (Supplementary Fig. 6c). To determine whether the increase in replication stress following WEE1 inhibition was due to an increase in DNA breakage, we used alkaline and neutral comet assays to measure the single-stranded DNA (ssDNA) breaks and double-stranded DNA (dsDNA) breaks, respectively. Results show that MK1775 treatment in RBM10-KO cells led to increased accumulation of both ssDNA and dsDNA breaks, with the increase in ssDNA breaks being more prominent (Fig. 5c,d and Supplementary Fig. 6d). Next, we utilized native BrdU staining to further confirm the increase in ssDNA in H1299^RBM10-KO^ cells upon MK1775 treatment (Fig. 5e,f). To address whether the accumulation of ssDNA upon WEE1 inhibition is due to ssDNA gaps arising during DNA replication, we used a modified DNA combing assay incorporating a digestion step with S1 nuclease^76^. Results revealed that labelled DNA tracks in MK1775-treated RBM10-KO cells showed high sensitivity to S1 nuclease digestion compared to MK1775-treated parental cells, suggesting WEE1 inhibition in RBM10-KO cells leads to accumulated ssDNA gaps during DNA replication (Fig 5g). Altogether, our results suggest that WEE1 inhibition exacerbates replication stress upon RBM10 loss and that replication stress underpins, at least partly, RBM10-WEE1 synthetic lethality.

**Figure 5:**
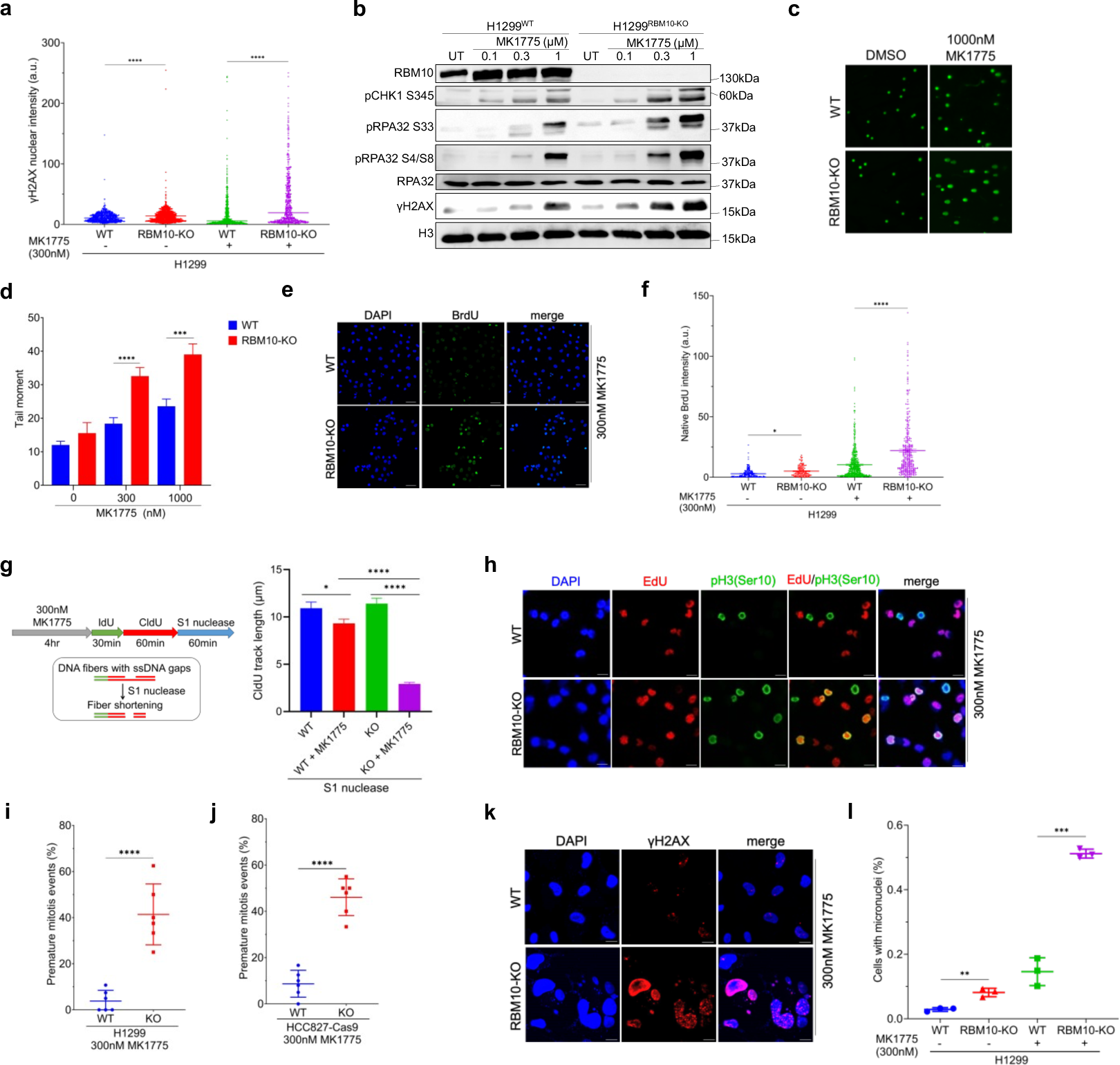
DNA damage and premature mitotic entry underpin RBM10-WEE1 synthetic lethality. **(a)** Quantification of γH2AX staining in H1299^WT^ and H1299^RBM10-KO^ cells treated with 300nM MK1775 for 24 hr. Data are presented as mean nuclear intensity ± SEM (n>590 cells) and representative of 3 independent experiments. Statistical significance was determined by Mann-Whitney test. *\*\*\*\*P<0.0001*. **(b)** Immunoblot analysis of DNA damage and replication stress markers in H1299^WT^ and H1299^RBM10-KO^ cells after MK1775 treatment. Cells were either left untreated (UT) or treated with the indicated concentrations of MK1775 for 24 hr followed by immunoblot analysis using the indicated antibodies. Histone H3 is used as a loading control. The positions of molecular weight markers are indicated to the right. **(c,d)** Alkaline comet assay in H1299^WT^ and H1299^RBM10-KO^ cells treated either with DMSO or the indicated concentrations of MK1775 for 24 hr. (c) Representative fluorescence microscopy images of alkaline comet assay. (d) Quantification of DNA damage represented by comet tail moment. Data are presented as mean tail moment ± SEM (n>150) and representative of 3 independent experiments. Statistical significance was determined by Mann-Whitney test. *\*\*\*P<0.001;****P<0.0001*. **(e,f)** Analysis of ssDNA levels using native BrdU staining in H1299^WT^ and H1299^RBM10-KO^ cells treated with 300nM MK1775 for 24 hr. (e) Representative immunofluorescence microscopy image for native BrdU staining. DAPI is used to stain nuclei. Scale bar, 50µm. (f) Quantification of native BrdU staining. Data are presented as mean nuclear intensity ± SEM (n>100 cells for untreated cells, n>300 for MK1775-treated cells) and representative of 3 independent experiments. Statistical significance was determined by Mann-Whitney test. *\*\*\*\*P<0.0001*. **(g)** H1299^WT^ and H1299^RBM10-KO^ cells were either untreated or treated with 300nM MK1775 for 4 hr followed by DNA combing analysis as indicated. The length of CldU+ region of IdU+ CldU+ replication tracks was measured following digestion with S1 nuclease for 1 hr. Data are presented as mean track length ± SEM (n>100). Statistical significance was determined by Mann-Whitney test. *\*P<0.05*;*\*\*\*\*P<0.0001*. **(h)** Representative immunofluorescence microscopy images showing premature mitotic entry in H1299^WT^ and H1299^RBM10-KO^ cells treated with MK1775. EdU incorporation and pH3(Ser10) were used to determine DNA synthesis and mitotic entry, respectively. DAPI is used to stain nuclei. Scale bar, 20µm. **(i,j)** Quantification of premature mitotic entry events in parental (WT) and H1299^RBM10-KO^ (i) and HCC827^RBM10-KO^ (j) cells. Premature mitosis events were calculated as the percentage of EdU-positive cells from the pH3(Ser10)-positive (mitotic) cells and presented as mean ± s.d. (n=6). Statistical significance was determined by unpaired t-test. *\*\*\*\*P< 0.0001*. **(k)** Representative immunofluorescence microscopy images of γH2AX and DAPI staining in H1299^WT^ and H1299^RBM10-KO^ cells treated with 300nM MK1775 for 24 hr. Scale bar, 10µm. **(l)** Quantification of micronuclei in H1299^WT^ and H1299^RBM10-KO^ cells treated with 300nM MK1775 for 24 hr. Data are presented as the average percentage of cells with micronuclei ± SEM (n=3). Statistical significance was determined by unpaired t-test. *\*\*\*\*P<0.0001*.

Next, we sought to determine the effect of WEE1 inhibition on cell cycle progression in RBM10-deficient cells. Towards this, parental and H1299^RBM10-KO^ cells were co-stained for pH3(Ser10), a marker of early mitosis, and EdU as a marker for S-phase following MK1775 treatment. Remarkably, unlike parental cells, a significant fraction (∼45%) of MK1775-treated H1299^RBM10-KO^ cells were co-stained with EdU and pH3(Ser10), suggesting that WEE1 inhibition leads to premature mitotic entry in RBM10-deficient cells (Fig. 5h,i). Similar results were also obtained in parental and HCC827^RBM10-KO^ cells (Fig. 5j). Consequently, we observed that MK1775-treated H1299^RBM10-KO^ cells exhibit high levels of micronucleation, extensive nuclear γH2AX staining and apoptosis (Fig. 5k,l and Supplementary Fig. 6e). Altogether, we concluded that the sensitivity of RBM10-deficient cells to MK1775 is mediated by increased replication stress and premature mitotic entry leading to mitotic catastrophe and cell death.

### MK1775 inhibits the growth of RBM10-deficient LUAD tumors *in vivo*

Given that the WEE1 inhibitor MK1775 is currently undergoing phase II clinical trials for treating several tumor types^77^, we sought to test the therapeutic potential of WEE1 inhibition against RBM10-deficient LUAD cells *in vivo*. Our results showed that MK1775 treatment prominently reduced the size of xenograft tumors derived from HCC827^RBM10-KO^ and H1299^RBM10-KO^ cells, but not their parental counterparts (Fig. 6a,b and Supplementary Fig. 7a,b). Consistent with the observed γH2AX phenotype *in vitro*, RBM10-deficient tumors exhibit increased levels of DNA damage in MK1775-treated mice (Fig. 6c,d and Supplementary Fig. 7c).

**Figure 6:**
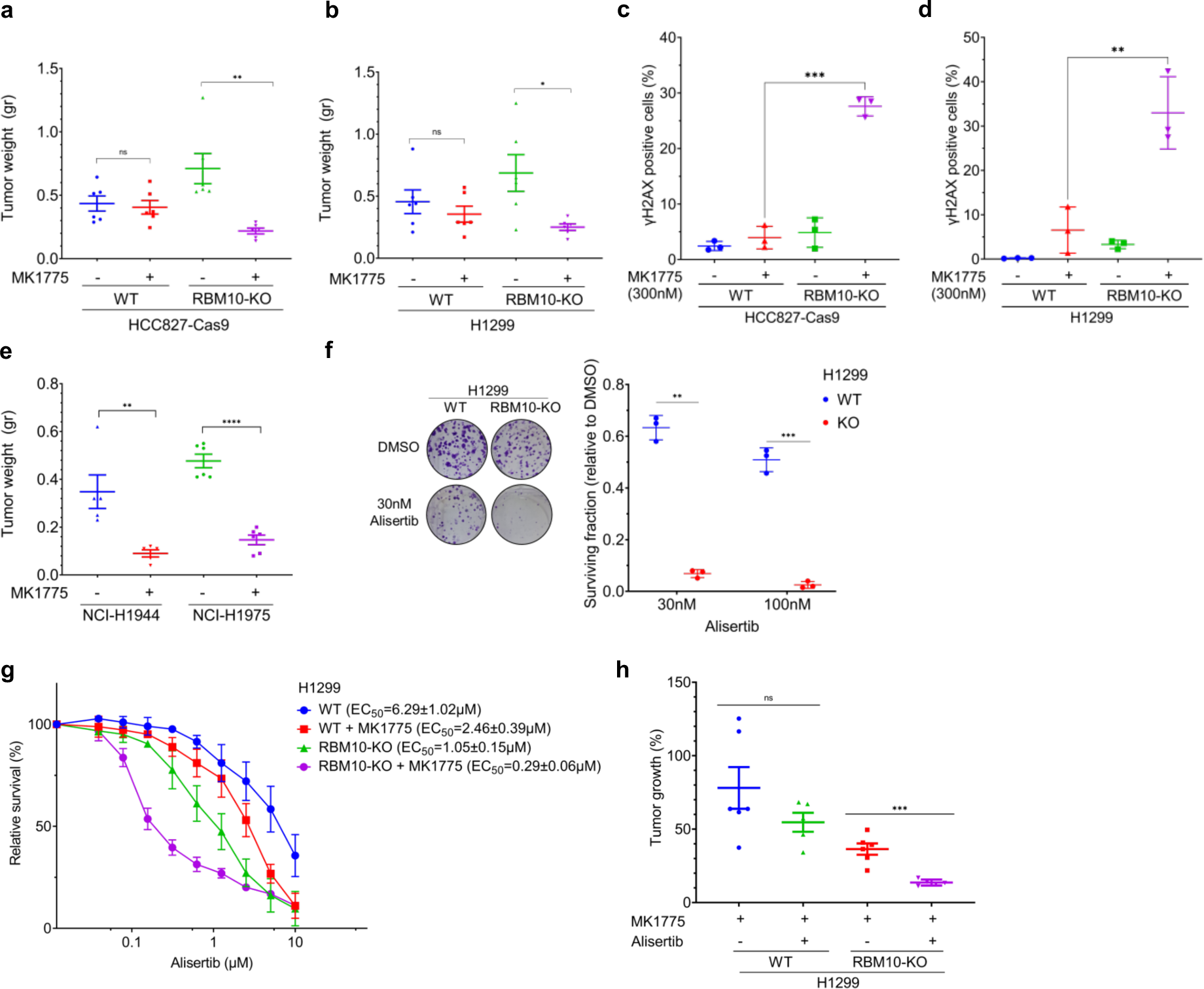
MK1775 inhibits the growth of RBM10-deficient LUAD cells *in vivo*. **(a,b)** Tumor weight of parental (WT) and HCC827-Cas9^RBM10-KO^ (a) and H1299^RBM10-KO^ (b) xenografts treated with either MK1775 or vehicle. MK1775 was administered once daily at 40mg/kg for 15 days. Results are shown as mean tumor weight ± SEM (n=6). Statistical significance was determined by unpaired t-test. *\*P<0.05;**P<0.01*. **(c,d)** Quantification of γH2AX immunostaining in xenograft tumor sections from parental and HCC827-Cas9^RBM10-KO^ (c) and H1299^RBM10-KO^ (d) tumors treated with either MK1775 or vehicle. Data are presented as the average percentage γH2AX positive cells ± SEM of 3 different tumor sections for each group. Statistical significance was determined by unpaired t-test. *\*\*P<0.01;***P<0.001.* **(e)** Tumor weight of NCI-H1944 and NCI-H1975 xenografts treated with either MK1775 or vehicle. MK1775 was administered once daily at 40mg/kg for 15 days. Results are shown as mean tumor weight ± SEM (n=5 for NCI-H1944, n=6 for NCI-H1975). Statistical significance was determined by unpaired t-test. *\*\*P<0.01;****P<0.00001*. **(f)** Clonogenic survival of H1299^WT^ and H1299^RBM10-KO^ cells treated with the indicated concentrations of alisertib. Left, representative images of plates stained with crystal violet. Right, quantification of clonogenic survival. Data are presented as mean ± s.d. (n=3). **(g)** Short-term cell viability assay and EC_50_ determination in H1299^WT^ and H1299^RBM10-KO^ cells treated with increasing concentrations of alisertib with or without treatment with 300nM MK1775. Data are presented as mean ± s.d. (n=3). **(h)** Tumor growth inhibition in H1299^WT^and H1299^RBM10-KO^ xenografts treated with MK1775 alone, MK1775 and alisertib combination, or vehicle. MK1775 alone or MK1775 and alisertib combination were administered once daily at 30mg/kg for 15 days. Results are shown as percentage tumor growth inhibition relative to vehicle ± SEM (n=6 for MK1775 alone and vehicle, n=5 for MK1775 + alisertib combination). Statistical significance was determined by unpaired t-test. *\*\*\*P<0.001*.

Collectively, these findings strongly suggest that RBM10 loss leads to increased sensitivity to WEE1 inhibition *in vivo*, independently of EGFR or TP53 status. To further substantiate the clinical relevance of targeting RBM10-deficient LUAD, we tested the anti-tumor effect of MK1775 against tumor xenografts derived from patient-derived LUAD cells, NCI-H1944 and NCI-H1975, harboring naturally occurring RBM10 cancer mutations. Results showed that WEE1 inhibition leads to remarkable tumor growth inhibition in RBM10-deficient NCI-H1975 and NCI-H1944 compared to RBM10-proficient cells (Fig. 6e). Altogether, our results demonstrate the therapeutic efficacy of MK1775 as a single-agent in eradicating LUAD tumors harboring RBM10 deleterious cancer mutations.

### Aurora kinase A inhibition exacerbates the anti-tumor activity of WEE1 inhibition in RBM10-deficient LUAD

Given that both WEE1 and Aurora A (AURKA) scored highly in RBM10 SL screen and both kinases are targeted by small molecule inhibitors undergoing clinical trials, we tested the efficacy of the combined Aurora A and WEE1 inhibition on the growth of RBM10-deficient LUAD. First, we observed that the Aurora A inhibitor, alisertib, had a profound effect on the clonogenic survival of RBM10 KO cells, concurrent with the CRISPR SL screen results (Fig. 6f and Supplementary Fig. 7d). Moreover, while alisertib alone led to a significant reduction in RBM10-deficient cell viability, combined MK1775 and alisertib treatment further exacerbated cell toxicity (Fig. 6g). Prompted by this, we tested the effect of the combined MK1775 and alisertib treatment on the growth of RBM10-deficient xenograft tumors. Results showed that the combined treatment leads to a profound and significant reduction in RBM10-deficient tumor size that is greater than the reduction upon MK1775 treatment alone (Fig. 6h). Therefore, we concluded that Aurora kinase A inhibition might be a viable therapeutic strategy to enhance the anti-tumor activity of WEE1 inhibition in RBM10-deficient LUAD tumors.

## Discussion

We present a model that underscores a new function of RBM10 in both DNA replication progression and replication stress response (Fig. 7). RBM10 interactome analysis revealed that it interacts with various replication fork components that are required for proper replication fork progression, as well as recovery from replication stress. Also, we show that RBM10 depletion leads to replication fork slowdown, which is accompanied by an increase in DNA damage. Our data are in line with a genome-wide siRNA screen that identified RNA processing factors, including RBM10, as replication stress response genes^56^. Moreover, previous studies showed that RBM10 is associated with human pre-replication complex^78^ and interacts with several proteins involved in DNA replication and repair in fission yeast^36^. In this regard, it would be of interest to determine whether other RNA splicing and processing factors are also implicated in DNA replication. In this study, we show that the RNA-binding protein RBM10 is recruited to active DNA replication forks in a PRIM1-dependent manner to fine-tune the levels of histone acetylation at active DNA replication fork. Given that PRIM1 knockdown alleviates RBM10 association with active replication forks, we propose a model that this association requires RNA molecules, presumably the RNA primers of Okazaki fragments. In this regard, we revealed that the RBM10 ZnF1 domain, known for its RNA-binding capability, mediates its association with replication fork. Interestingly, this ZnF1 domain is found in various proteins including MDM2, NEIL3, ZRANB3, and FUS that have been previously implicated in regulating DNA replication progression and stress response^79–82^. Future investigations will be necessary to determine whether the association of these proteins with DNA replication forks is mediated by the RNA primers of Okazaki fragments. Functionally, we demonstrate that RBM10 counteracts the formation of R-loops in S-phase cells, concurrently facilitating the association of HDAC1 with active replication forks to deacetylate H4K16. These findings align with recent studies that described a role of HDAC1/2 in chromatin remodeling at active replication forks to maintain normal replication fork progression and replication fork stability^64–66^. Interestingly, it was shown that HDAC1/2 decrease the levels of the transcriptionally active H4K16 acetylation at replication forks, thereby reducing the levels of R-loops and mitigating transcription-induced replication stress^66^. Further work will be required to determine the impact of RBM10 depletion on the acetylation levels of other histone and non-histone residues that are implicated in DNA replication regulation. While our data suggest that RBM10 function in DNA replication is distinct from its splicing activity (Fig. 3g), we cannot exclude the possibility that RBM10 may regulate the splicing or expression of genes involved in regulating replication fork progression.

**Figure 7:**
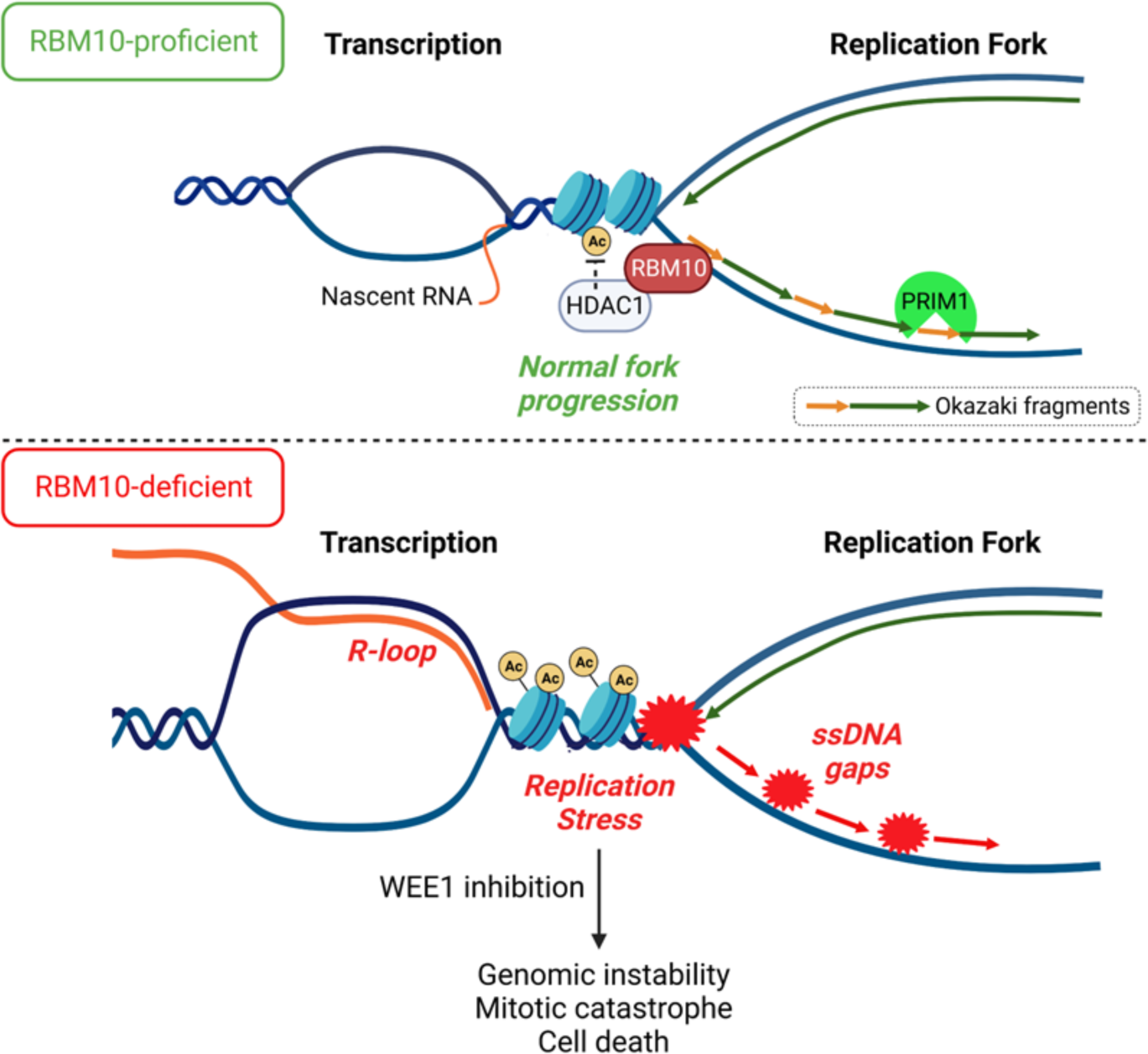
Model depicting the novel role of RBM10 in DNA replication and replication stress response. (Top) RBM10 association with active DNA replication forks is dependent on PRIM1, which synthetizes the RNA primer of Okazaki fragments. RBM10 promotes the recruitment of HDAC1 to ongoing and stressed replication forks and ensures the deacetylation of H4K16, thereby limiting R-loop formation and maintaining fork stability. (Bottom) In RBM10-deficient cells, defective HDAC1 recruitment and H4K16 deacetylation at stressed replication forks contributes to R-loop accumulation, fork destabilization, and ssDNA gap formation leading to replication stress. High levels of replication stress render RBM10-deficient tumor cells sensitive to WEE1 kinase inhibition, leading to the accumulation of DNA damage and mitotic catastrophe resulting in tumor cell death. Created with BioRender.com.

In this study, we performed a genome-wide CRISPR-Cas9 knockout screen employing isogenic LUAD cells carrying RBM10 cancer mutation, a type of nonsense mutation occurring with frequencies ranging from 9% to 25% in LUAD. We identified ∼250 RBM10 SL genes, such as WEE1 and Aurora A kinases, which can be therapeutically exploited for eradicating RBM10-deficient LUAD tumors. Indeed, pharmacological inhibition of WEE1 using MK1775 markedly reduces RBM10-deficient cell proliferation and tumor growth. Given that MK1775 is currently in phase II clinical trials for the treatment of several types of tumors, we propose that it can be also used against RBM10-deficient LUAD with immediate clinical applicability. Interestingly, the sensitivity of RBM10-deficient cells to WEE1 inhibition is further exacerbated by the combined treatment with alisertib, an Aurora Kinase A inhibitor. This synergistic effect between these two inhibitors is in agreement with a previous study showing that simultaneous inhibition of Aurora A and WEE1 prompts cells to traverse the G2–M checkpoint in the presence of DNA damage and impaired spindle assembly, culminating in mitotic catastrophe. Consequently, it leads to a synergic anti-tumor effect in p53-deficient squamous cell carcinoma of the head and neck^83^. Together, these observations provide a basis for testing this combination for clinical treatment of RBM10-deficient LUAD patients, thus increasing therapeutic benefits and minimizing the risk for developing resistance.

Mechanistically, RBM10-WEE1 synthetic lethality is mediated, at least in part, by increased replication stress leading to premature mitotic entry and cell death. This finding supports the notion that the sensitivity of RBM10-deficient cells to WEE1 inhibition is dependent on the emerging role of RBM10 in DNA replication. Our data are therefore in agreement with previous studies showing synergistic interaction between WEE1 inhibition and replication stress-inducing agents^84–87^.

Intriguingly, a previous report showed that RBM10 deficiency in LUAD is associated with enriched immune pathways, elevated tumor mutational burden, and increased human leukocyte antigens (HLA) expression, suggesting that RBM10 deficiency may enhance anti-tumor immunity in LUAD^88^. In agreement with this, our RBM10 transcriptome in HCC827 and H1299 cell lines showed that RBM10 depletion leads to increased expression of proinflammatory cytokines such as IL1R2, TNFRSF1B, BMP2, IL7R and CCL2. Further work will be vital to determine whether WEE1 inhibition in RBM10-deficient tumors further amplifies the upregulation of genes associated with immune pathways, which might enhance the response to immune therapies.

Unbiased genome-wide CRISPR-Cas9 SL screens revolutionized the discovery of novel SL interactions that can be exploited for developing new cancer therapies^89,90^. Our CRISPR-Cas9 knockout screen revealed ∼250 RBM10 SL genes involved in various cellular pathways, such as RNA splicing and cell cycle. These genetic interactions advance our understanding of the physiological functions of RBM10 and provide a repertoire of novel targets that can be harnessed to formulate new strategies for targeting RBM10-deficiency. Interestingly, RBM10 is frequently mutated in various human cancers such as endometrial (8%) and urothelial (5%) carcinomas^58^. Therefore, our findings might be of clinical interest to other types of human cancer harboring RBM10 mutations.

## Acknowledgments

We thank Avigail Yampolsky and Yarin Attar for helping in validating the RBM10 CRISPR-Cas9 screen results, Dvir Aran for his help in analyzing RBM10 mutations in cancer, Yossi Shiloh and Yehuda Assaraf for their insightful comments on the project. We also thank Nadav Sharon, Assaf Bester, and Juan Valcárcel for generously sharing reagents and plasmids. We thank Janette Zavin for histological processing. We thank our lab members for critical reading of the manuscript. We thank bioRENDER (biorender.com) for assisting in drawing the model and the schematics presented in the figures. Research in the Ayoub lab is supported by grants from the Israel Science Foundation (2511/19) and Israel Cancer Research Fund (22-110-PG). Research in the Simon lab is supported by grants from the Israel Science Foundation (1283/21) and the Binational Science Foundation (2019688 and 2021085). F.E.M. and E.R.A. are supported by Clore fellowship. A.B. is supported by Joan and Irwin Jacobs fellowship. N.A. is supported by the Neubauer Family foundation.

## Author Contributions

F.E.M. wrote the original draft and performed all the experiments and analysis described in this manuscript except the ones indicated below. E.R.A. cloned flag-MCM5 expression plasmid and helped in setting up the CRISPR-Cas9 screen and proofreading the manuscript. J.K. performed the experiments described in Figure 2a, 2b, 5g and Supplementary Figure 2e, 5c. A.B. performed the experiments described in Supplementary Figure 2d, 3d and helped in characterizing RBM10 splicing activity. I.S. helped in analyzing the data related to RBM10 role in DNA replication progression and editing the manuscript. N.A. conceived and supervised the study.

## Corresponding Author

Correspondence to Nabieh Ayoub.

## Methods

### Cell lines and cell culture

HCC827, NCI-H1299, NCI-H1944, and NCI-H1975 cell lines were obtained from ATCC and grown at 37°C and 5% CO2 and cultured in RPMI-1640 medium (Gibco) supplemented with 10% heat-inactivated fetal bovine serum (Gibco), 2mM L-glutamine (Gibco), and 100unit/mL penicillin and 100μg/mL streptomycin (Gibco). HEK293T cell line was obtained from ATCC and grown at 37°C and 5% CO2 and cultured in Dulbecco’s Modified Eagle Medium (DMEM) (Gibco) supplemented with 10% heat-inactivated fetal bovine serum (Gibco), 2mM L-glutamine (Gibco), and 100unit/mL penicillin and 100μg/mL streptomycin (Gibco).

### Lentiviral transduction

Lentiviral particles were generated by co-transfecting HEK293T cells plated in 10cm plates with 1.64pmol target vector, together with 1.3pmol psPAX2 (Addgene #12260) and 0.72pmol pMD2.G (Addgene #12259) using 3:1 μg PEI to μg DNA ratio. Media containing the viral particles were collected 48 hr post-transfection and filtered with 0.45 μm filters. The indicated cell lines were transduced with the lentiviral particles in the presence of 10μg/ml polybrene (Sigma-Aldrich H9268). At 48 hr post-infection, transduced cells were selected using the appropriate antibiotics.

### Generation of Flag-Cas9 expressing cells and RBM10 knockout cell lines

To generate HCC827 cell line constitutively expressing Cas9, cells were transduced with Lenti-Cas9-2A-Blast (Addgene #73310) lentiviral particles followed by selection with 10μg/ml blasticidin (Invivogen #ant-bl) for 72 hr. A single clone stably expressing Flag-Cas9 fusion was selected and validated by western blot and immunofluorescence. To generate isogenic RBM10 knockout in HCC827 cells expressing flag-Cas9 (HCC827-Cas9), guide RNAs (gRNA) were designed and cloned into pSPgRNA plasmid (Addgene #47108) using BbsI restriction enzyme. HCC827-Cas9 cells were transfected with RBM10 gRNA1 (HCC827-Cas9^RBM10-KO^ and naïve HCC827^RBM10-KO3^) or gRNA2 (HCC827-Cas9^RBM10-KO2^) followed by clonal isolation. Single clones were screened for RBM10 knockout using western blot and validated by immunofluorescence and sequencing. The sequences of RBM10 gRNAs are included in Supplementary Table 3. To generate RBM10 knockout in naïve HCC827 and NCI-H1299 cells, RBM10 gRNA1 was cloned into pSpCas9(BB)-2A-GFP (PX458) (Addgene #48138) and transfected to cells. At 24 hr after transfection, GFP-positive cells were sorted using BD LSRFortessa™ cell analyzer (BD Biosciences) and plated in 96-well plates at a dilution of one cell per well. Single clones were screened for RBM10 knockout using western blot and validated by immunofluorescence and sequencing.

### Genome-wide CRISPR-Cas9 synthetic lethal screen

240 million cells of HCC827-Cas9^WT^ and HCC827-Cas9^RBM10-KO^ were transduced in triplicates with TKOv1 sgRNA library, which contains 91,320 sgRNAs targeting 17,232 protein-coding genes^52^, at MOI=0.3 and an average of 500-fold coverage of the library. 24 hr post-transduction, cells were selected with 2 μg/mL puromycin for 2 days. Following selection, 80 × 10^7^ cells were harvested as day 0 samples (T0) and the remaining cells were plated and passaged every 5 days. Cells were then collected at day 15 (T15) for isolation of genomic DNA (gDNA) together with the T0 samples. Genomic DNA extraction was performed with Blood & Cell Culture DNA Maxi Kit (Qiagen) according to the manufacturer’s instructions. gRNA inserts were amplified using two rounds of PCR reactions. First, a 600bp region of the lentiviral library vectors containing the sgRNA region was amplified using ordinary PCR with Q5 high-fidelity DNA polymerase for 25 cycles. A total of 120μg gDNA was used as template for each sample (60 reactions: 2μg gDNA/50μl reaction). Second, the amplified PCR products were subjected to another PCR reaction with Illumina next-generation sequencing adaptors for 10 cycles and 200bp product was amplified. The resulting amplicons were pooled and gel purified using the NucleoSpin Gel and PCR Clean-up (Macherey-Nagel). Samples were sequenced on a MiSeq platform with ∼10 million reads per sample (Illumina). Demultiplexing and mapping of sequencing reads to the gRNA library was performed using MAGeCK^54^. Fold-change for each gRNA was calculated in each replicate versus the T0 sample. 3 analysis methods were used to predict RBM10 synthetic lethal genes: (1) Bioinformatic and Bayesian Analysis of Gene Essentiality (BAGEL) algorithm^53^ was devised to calculate the essentiality score (Bayes factor – BF) of each gene in parental (WT) and RBM10-KO cells. RBM10 SL genes were defined as genes with BF>5 (i.e. essential) in RBM10-KO cells and BF<2 (i.e. non-essential) in WT cells. (2) CRISPR Counts Analysis (CCA)^55^ was used to calculate synthetic lethality score for each gene and genes with score > 0.8 were considered candidate RBM10 SL genes. (3) MAGeCK was used to calculate log fold-change (LFC) for each gene and genes with LFC<- 1 in RBM10-KO cells with LFC difference > 1 between WT and RBM10-KO were considered RBM10 SL genes. Pathway enrichment analysis and gene ontology were conducted using ShinyGO^91^. STRING interaction network was performed using string-db.org.

### Conditional short hairpin RNA (shRNA) knockdown

shRNA oligonucleotides directed against the indicated genes were annealed and cloned into Tet-pLKO-puro lentiviral vector (Addgene #21915) digested with *Eco*RI and *Age*I. Lentiviral particles were generated as described above and used to transduce parental and RBM10-KO HCC827-Cas9 cells, followed by selection with 1μg/ml puromycin (Invivogen #ant-pr) for 3 days. Cells were maintained in complete medium in the presence of 0.6μg/ml puromycin. To induce shRNA-mediated gene expression knockdown, cells were cultured in the presence of 0.2μg/ml doxycycline (DOX) for 48 hr prior to clonogenic survival assays. Gene expression knockdown was validated by RT-qPCR. The shRNA and qPCR primer sequences used in this study are included in Supplementary Table 4.

### Generation of expression vectors

To generate EGFP-RBM10 expression vectors, RBM10 was amplified from pDest26-RBM10-wt plasmid^32^ (gift from Prof. Juan Valcárcel) and cloned into pEGFP-C1 expression plasmid. To generate RBM10 deletion mutants, all-round PCR was used to delete the relevant regions. To constitutively restore RBM10 expression in RBM10-deficient cells, full-length RBM10 and RBM10 deletion mutants were subcloned to Lenti-Cas9-2A-Blast plasmid followed by lentivirus generation and transduction. To generate Flag-MCM5 expression vector, MCM5 was cloned from cDNA into p3X-Flag-CMV10 expression vector. Primer sequences used for cloning and mutagenesis are included in Supplementary Table 4.

### Transfections

Cell transfections with plasmid DNA or siRNA were performed using Polyethylenimine (PEI) and Lipofectamine RNAiMax, respectively, following the manufacturer’s instructions. siRNAs used in this study are: RBM10 siRNA #1 (ThermoFisher Scientific HSS112075, 5′-CAAACGCCGAGAGAAGUGCU-3′); RBM10 siRNA #2 (ThermoFisher Scientific HSS112075, 5’-TTTGCCAAGGGTTCTAAGAG-3’); PRIM1 siRNA (Euphera Biotech EHU105881); Stealth RNAi negative control (Invitrogen).

### RNA isolation, RT-PCR and qPCR

Total RNA was isolated from cells using the TRIzol reagent according to the manufacturer’s instructions (Ambion) and treated with RQ1 DNase (M6101, Promega). 1μg RNA was used for cDNA synthesis using the qScript cDNA Synthesis Kit (Quanta) with random primers. RT-PCR for alternative splicing assay of NUMB and EIF4H genes was performed by standard PCR using Red Load Taq Master (Jena Bioscience) followed by gel electrophoresis in agarose gel stained with ethidium bromide. Band intensities were quantified using GelDoc software. Real-time qPCR for measuring mRNA levels was performed using Step-One-Plus real-time PCR System (Applied Biosystems) and the Fast SYBR Green Master mix (Applied Biosystems) with three technical repeats for each PCR. Data analysis and quantification were performed using StepOne software V2.2 supplied by Applied Biosystems. Primer sequences used for RT-PCR and qPCR are included in Supplementary Table 4.

### Small molecule inhibitors and drugs

MK1775 (A5755), Alisertib (MLN8237) (A4110), and hydroxyurea (B2102) were purchased from APExBio. Caffeine was purchased from Sigma-Aldrich (C8961).

### Immunoblotting

Whole-cell protein extracts were prepared using hot lysis buffer (1%SDS, 5mM EDTA, 50mM Tris, pH7.5) supplemented with protease inhibitor mixture (Calbiochem). Protein samples were diluted and boiled in 5X protein loading buffer (10% SDS, 500mM DTT, 50%Glycerol, 250mM Tris-HCL and 0.5%bromophenol blue dye) and separated on an SDS-polyacrylamide gel. Proteins were transferred to a PVDF membrane (Millipore) and immunoblotted with the indicated antibodies. The following primary antibodies were used for immunoblotting: RBM10 (Sigma-Aldrich cat.no. HPA034972, 1:10,000), FLAG-tag (Sigma-Aldrich cat.no. F7425, 1:4,000), β-actin (Sigma-Aldrich cat.no. A5441, 1:10,000), α-Tubulin (Santa Cruz Bio cat.no. sc-5286, 1:1,000), pRPA32-S4/S8 (Bethyl cat.no. A300-245A, 1:1,000), pRPA32-S33 (Bethyl cat.no. A300-246A, 1:1,000), RPA32 (Abcam cat.no. ab2175, 1:500), γH2AX (Cell Signaling Technology cat.no. 2577, 1:1000), pChk1-S345 (Cell Signaling Technology cat.no. 2348, 1:1,000), Histone H3 (Abcam cat.no. ab1791, 1:30,000), Rad51 (Santa Cruz Bio cat.no. sc-8349, 1:1,000), PARP1 (Enzo Life Sciences cat. no. ALX-210-895, 1:2,000), HDAC1 (GeneTex cat. no. GTX100513, 1:2,000), PCNA (Santa Cruz Bio cat. no. sc-56, 1:1,000), GAPDH (Abcam cat.no. ab8245, 1:1,000), GFP (Santa Cruz Bio cat.no. sc-9996, 1:1,000). The following secondary antibodies were used: anti-rabbit IgG-HRP (Jackson ImmunoResearch cat.no. 111–035-003, 1:20,000), anti-mouse IgG-HRP (Amersham cat.no. NXA931, 1:10,000). Immunoblot intensity quantification was performed using ImageJ.

### Biochemical fractionation

Cells were lysed with Buffer A for 5 min at 4°C. Cell lysates were centrifuged at 1500× g for 5 min at 4°C and the supernatant (cytoplasmic fraction) was removed. Then, the pellet was incubated with Buffer B (3mM EDTA, 0.2mM EGTA, 1mM DTT, PMSF, and protease inhibitor mixture) for 10 min on ice followed by centrifugation at 1700× g for 5 min at 4°C to extract nuclear-soluble fraction. To prepare chromatin-bound fraction, pellet was resuspended with hot lysis buffer (1%SDS, 5mM EDTA, 50mM Tris, pH7.5), boiled for 15 min, and sonicated with two 15 sec pulses of 35% amplitude, centrifuged at maximum speed for 20 min at 12°C, and the supernatant was recovered. Fractions were subjected to immunoblot analysis with the indicated antiabodies. For chromatin-bound fractionation with RNaseA, pellet was first resuspended in Buffer A containing 5 µg RNase A and incubated for 2 hr at room temperature followed by extraction with hot lysis buffer.

### GFP-trap

HEK293T cells expressing either EGFP only or EGFP-RBM10 plasmids were subjected to GFP-TRAP assay as previously described^92^. Where indicated, cells were also co-transfected with Flag-MCM5 plasmid. Briefly, EGFP-RBM10 or EGFP-only expressing cells were lysed in RIPA buffer (150mM NaCl, 50mM Tris pH 7.4, 1% NP40, 0.5% deoxycholate, 0.1% SDS) supplemented with protease inhibitor cocktail and then diluted 1:4 with binding buffer (150mM NaCl, 50mM Tris pH 7.4). 2 milligram of cell lysate was incubated for 4 hr at 4°C with 15μl GFP-TRAP beads followed by western blot analysis. For GFP-trap followed by mass spectrometry (MS), HEK293T cells were transfected with EGFP-RBM10, lysed and subjected to GFP-trap as described above. Beads were washed 5 times in PBSx1 followed by elution and trypsin digestion. Digested peptides were then analyzed by mass spectrometry (MS) at the Smoler Proteomics Center in the department of Biology in the Technion. The samples were analyzed by LC-MS/MS using Q-Exactive plus mass spectrometer (Thermo Scientific), coupled to Easy nano LC-1000 capillary UHPLC (Thermo Scientific). Protein identification and intensity analysis was performed using maxquant^93^, searching against the human section of the Uniprot database. RBM10-interacting proteins enrichment analysis and gene ontology was conducted using ShinyGO^91^.

### Immunoprecipitation

For endogenous RBM10 immunoprecipitation, H1299 cells were harvested and lysed in RIPA buffer (150mM NaCl, 50mM Tris pH 7.4, 1% NP40, 0.5% deoxycholate, 0.1% SDS) supplemented with protease inhibitor cocktail. Immunoprecipitations were performed using 0.5μg IgG or RBM10 antibody and protein A magnetic beads (GenScript #L00273) incubated with 2mg lysate overnight at 4°C. Immunoprecipitated samples were subjected to immunoblot analysis using the indicated antibodies.

### Immunofluorescence

Cells were seeded onto glass coverslips and grown for 24 hr followed by drug treatments as indicated in the figure legends. Cells were fixed with 4% (wt/vol) paraformaldehyde for 10 min, permeabilized with 0.5% Triton X-100 in PBS for 10 min, blocked with blocking buffer (4% (wt/vol) BSA, 0.15% Tween 20 and 0.15% Triton X-100 in PBS) for 1 hr at room temperature and incubated with the indicated antibodies for 1 hr at 37°C. Excess antibody was washed three times with wash buffer (0.15% Tween 20 and 0.15% Triton X-100 in PBS × 1), and cells were stained with secondary antibodies for 1 hr at room temperature in the dark, and then washed as above. Coverslips were mounted onto glass slides using VECTASHIELD® Antifade Mounting Medium with DAPI (Vectorlabs). For immunofluorescence staining of γH2AX, pRPA32-S33, and BrdU, cells were pre-extracted with 0.25% Triton X-100 in PBS for 15 min on ice prior to fixation. For EdU incorporation, cells were pulsed with 10μM EdU (5-ethynyl-2-deoxyuridine, Invitrogen A10044) for 30 min at 37°C, 5% CO2. EdU click reactions were performed after cell permeabilization by incubating cells with EdU staining buffer containing 2mM CuSO4, 100mM ascorbic acid and 1μM AlexaFluor 647 azide (Invitrogen A10277) in PBS for 30 min at room temperature. Images were acquired using inverted Zeiss LSM-700 or LSM-710 confocal microscopes. Image analysis was performed using ImageJ software. The following primary antibodies were used for immunofluorescence: BrdU FITC (BD Biosciences, cat.no. 51-23614 1:1,000), RBM10 (Sigma-Aldrich cat.no. HPA034972, 1:2,000), pRPA32-S33 (Bethyl cat.no. A300-246A, 1:500), γH2AX (Millipore cat.no. 05-636, 1:2,500), FLAG-tag (Sigma-Aldrich cat.no. F7425, 1:1,000), pH3(Ser10) (Cell Signaling Technology cat.no. 9706, 1:1,000), Anti-DNA-RNA Hybrid Antibody, clone S9.6 (Sigma-Aldritch cat.no. MABE1095, 1:1,000). The following secondary antibodies were used: anti-rabbit IgG Alexa Fluor^®^488 (Invitrogen cat.no. A-21206, 1:500), anti-mouse IgG Alexa Fluor^®^488 (Invitrogen cat.no. A-21202, 1:500), anti-rabbit Alexa Fluor^®^568 (Invitrogen cat.no. A10042, 1:500), anti-mouse Alexa Fluor^®^568 (Invitrogen cat.no. A10037, 1:500), anti-rabbit Alexa Fluor^®^647 (Invitrogen cat.no. A-31573, 1:500), anti-mouse IgG Alexa Fluor^®^647 (Invitrogen cat.no. A32787, 1:500).

### Proximity ligation assay (PLA)

Proximity ligation assay (PLA) was performed using the Duolink in situ red starter kit (Sigma-Aldrich) following the manufacturer’s instructions. Rabbit anti-RBM10 (Sigma-Aldrich HPA034972, 1:2,000) and mouse anti-PCNA (Santa Cruz Bio cat. no. sc-56, 1:250) antibodies were incubated for 1 hr at 37°C prior to PLA analysis. PLA assay between RBM10 and EdU-labelled nascent DNA was performed as previously described^94^. Briefly, cells were pulse labelled with 100μM EdU for 10 min at 37°C, followed by pre-extraction in 0.25% Triton X-100 in PBS for 10 min on ice and fixation in 4% (wt/vol) paraformaldehyde. Cells were incubated in EdU click reaction buffer containing 2mM CuSO4, 100mM ascorbic acid and 10μM biotin-azide (Invitrogen B10184) in PBS for 1 hr at room temperature. Cells were incubated in blocking buffer for 1 hr at room temperature followed by incubation with rabbit anti-RBM10 (Sigma-Aldrich HPA034972, 1:2000), rabbit anti-HDAC1 (GeneTex GTX100513, 1:2,000), Rabbit anti-H4K16ac (Abcam ab109463, 1:2,000), rabbit anti-53BP1 (Novus biological NB100-305, 1:1,000) and mouse anti-biotin (Jackson ImmunoResearch cat.no. 200-002-211, 1:2000) antibodies for 1 hr at 37°C. Next, PLA assay was performed using the Duolink in situ red starter kit for experiments described in Fig. 3c,d (Sigma-Aldrich) or NaveniFlex Cell MR Atto647N kit (Navinci) following the manufacturer’s instructions. Images were acquired using inverted Zeiss LSM-700 confocal microscope. Image analysis was performed using ImageJ software. PLA intensity was calculated as the product of number of spots and the mean intensity of the spots per nucleus.

### Cell synchronization

To synchronize cells at G1/S border using double-thymidine block, cells were treated with 2 mM thymidine overnight, released for 10 h and treated again with thymidine overnight. Cells were released into growth medium containing nocodazole (120 ng/ml) to enrich mitotic cells. Cell cycle profiles were analyzed by Cytek Aurora^©^ after staining with propidium iodide (Sigma-Aldritch).

### Alkaline and neutral comet assays

Alkaline and neutral comet assay was performed using the CometAssay Kit (Trevigen, 4250-050-K) according to the manufacturer’s instructions. Briefly, 1 × 10^5^ cells/mL were mixed with molten LMAgarose (at 37°C) at a ratio of 1:10 (v/v), and 70 μL of this mixture was transferred onto a CometSlide. Slides were placed flat at 4°C in the dark for 30 minutes. Cells were then lysed and subjected to alkaline or neutral electrophoresis with voltage of 1V/cm. DNA was stained with SYBR green and slides were imaged using ZEISS LSM700 confocal microscope. Comet tail moment was scored using CometScore 2.0 software.

### Short-term growth delay assay

For determining drug sensitivity, cells were seeded in 96-well plates in duplicates at a density of 2,000-5,000 cells per well. 24 hr post-seeding, drugs were added at the indicated concentrations. Cell viability was measured 72 hr after drug treatment using the CellTiter 96® AQueous One Solution Cell Proliferation Assay (Promega) following the manufacturer’s protocol, and absorbance was measured using Epoch Microplate Spectrophotometer (BioTek). Cell viability was normalized to the viability of untreated cells.

### Clonogenic survival assays

For colony survival assay, cells seeded onto 6-well plates (500 cells per well) and left untreated or treated with the indicated doses of drugs. After 14 days, cells were fixed in methanol and stained with 0.25% crystal violet. Individual colonies were counted, and relative survival was calculated by normalizing to the number of colonies in control wells.

### Annexin-V apoptosis assay

Apoptosis was assessed by annexin V-FITC kit (Invitrogen) according to the manufacturer’s instructions. Parental (WT) and RBM10-KO cells were treated with the indicated concentrations of MK1775 for 24 hr prior to analysis. Samples (10,000 cells) were analyzed using Cytek Aurora^©^. Results were calculated as the percentage of positive annexin V-FITC cells out of total cells counted.

### Endogenous homologous recombination assay

Homologous recombination repair assay was performed using Cas9-mediated knock-in of green fluorescent mClover into the first exon of the LMNA gene, as previously described^69^. Briefly, cells were plated in 6-well plates and co-transfected with 1.6 μg pX330-LMNA-gRNA1 plasmid containing Cas9 and gRNA against LMNA exon 1 and 0.4 μg pCR2.1-CloverLamin plasmid containing HR donor sequence. In addition, 0.4 μg pDsRed-Monomer-C1 was included per transfection as a transfection control unless otherwise indicated in the figure legend. 12−16 h post transfection, culture medium was renewed and where indicated, Caffeine (4 mM) was added. Seventy-two hours post-transfection, cells were collected and analyzed by Cytek Aurora^©^ . HR efficiency is percentage mClover-expressing cells from DsRed-Monomer positive cells.

### DNA combing assays

DNA combing analysis was performed as described^95^. Briefly, unsynchronized cells were pulse labelled with 25uM Idu for 30 min, followed by 600μM CldU for another 30 min, followed by 1.5 hr growth without any label. For combing analysis with hydroxyurea (HU), cells were pulse labelled with 25uM Idu for 30 min, followed by incubation with 4mM HU for 2 hr, followed by pulse labelling 600μM CldU for another 30 min. Cells were harvested into 2% agarose plugs, incubated for 48 hr in a digestion buffer (1% N Laurouylsarcosine, 0.2% Na Deoxycholate, 10mM Tris-HCl pH 7.5, 100mM EDTA, and 1mg/ml proteinase K). Next, TE 50X was added overnight, and the plugs were dissolved into 50mM MES pH6 with 100mM NaCl. The DNA was combed on Genomic vision combing slides, denatured and stained with the following primary antibodies: Rat anti-BrdU (Abcam #ab6326, 1:200), mouse anti-BrdU (BD Biosciences #347580, 1:20), and anti-ssDNA antibody (DSHB #AB10805144, 1:20). The following secondary antibodies were used: Goat anti-mouse Alexa Fluor^®^647 (Jackson Immunoresearch #115-605-166, 1:400), goat anti-rat Alexa Fluor^®^594 (ThermoFisher Scientific #A11007, 1:400), and anti-mouse IgG2a Alexa Fluor^®^488 (ThermoFisher Scientific #A21131, 1:500). The slides were photographed with a fluorescent microscope and analyzed using Olympus CellSens program. For DNA combing analysis with S1 nuclease digestion, cells were treated as indicated and then pulse labelled with 25uM Idu for 30 min, followed by 600μM CldU for 60 min. Subsequently, cells were permeabilized (10 mM PIPES, pH 6.8, 0.1 M NaCl, 0.3 M sucrose, 3 mM MgCl2, EDTA-free Protease Inhibitor Cocktail (Roche)) at room temperature for 10 min, followed by incubation with S1 nuclease (20 U/ml) (Promega M5761) in S1 buffer for 60 min at 37 °C

### RNA sequencing

Three biological replicates of RNA samples were purified from parental and RBM10-KO HCC827 and H1299 cells, or HCC827 cells transfected with control or RBM10 siRNA#1. RNA sequencing libraries were prepared using TruSeq mRNA library preparation kit. Sequencing was performed at The Crown Genomics institute of the Nancy and Stephen Grand Israel National Center for Personalized Medicine,Weizmann Institute of Science using a NovaSeq 6000 system with S1 flow cell to obtain 150 bp paired-end reads. Average read depth was 60 million reads per sample. The quality of the raw FASTQ files was assessed using FastQC software (http://www.bioinformatics.babraham.ac.uk/projects/fastqc/). For differential gene expression analysis, raw sequencing reads were aligned to GENCODE GRCh38 genome assembly using Salmon package^96^ and differential gene analysis was performed in R using the DESeq2 package^97^. Pathway enrichment analysis and gene ontology was conducted using ShinyGO^91^.

### Mouse xenograft model

Animal studies and protocols were approved by the Committee on the Ethics of Animal Experiments of the Technion, Israel Institute of Technology (IL-069-07-22). Immunocompromised NOD.CB17-Prkdcscid/NcrCrl (NOD SCID) mice were purchased from Envigo. Mouse model of LUAD cell lines subcutaneous xenografts were established by transplanting cells into 6-week-old female NOD SCID mice. Mice were randomly assigned into control and treatment groups. Prior to initiating any experiments, mice were allowed one week acclimation to housing conditions at the Technion Animal Facility. All mice were housed in a strict pathogen-free environment. 2.5 × 10^6^ of the indicated cells were resuspended in 100μl PBS and mixed 1:1 with Matrigel (High Concentration – Corning) and injected subcutaneously into the right flank of 6 weeks-old female NOD SCID mice. Tumors were measured using digital calipers. Tumor volumes were calculated using the formula: 0.5 × length × width^2^. When tumors reached ∼100mm^3^, mice were randomly assigned to treatment groups (n = 6 mice for each group, unless otherwise indicated). Each group received either control (vehicle), MK1775 (40mg/kg, p.o.), or MK1775 + alisertib (30mg/kg each, p.o.) once daily for 15 days. Mice were euthanized when the tumors of control mice reached ∼1500mm^3^ and tumors were excised, weighed, and photographed.

### Immunohistochemistry

Tumors excised from mice were fixed in 4% formalin at 4°C overnight, washed three times with PBS, and immersed in 30% (wt/vol) sucrose solution at 4°C overnight. Then, tumors were embedded in optimal cutting temperature compound (TissueTek) and frozen at −20°C and. 10μm cryosections were prepared and permeabilized using 0.5% Triton X-100 in PBS followed by immunofluorescence staining as described above, except primary antibody incubation with γH2AX antibody (Millipore cat.no. 05-636,1:500) were performed at 4°C overnight.

### Data availability

Raw next-generation sequencing data for RBM10 SL CRISPR-Cas9 screen have been deposited in ArrayExpress and are available through accession number E-MTAB-13626. RNA-seq sequencing data generated in this study have been deposited in ArrayExpress and are available through accession number E-MTAB-13617. The mass spectrometry proteomics data have been deposited to the ProteomeXchange Consortium via the PRIDE repository with the dataset identifier PXD040109. All data supporting the findings of this study are available from the corresponding author upon request.

