## Supplementary information for "Harnessing DNA replication stress to target RBM10 deficiency in lung adenocarcinoma"

### Supplementary Figures:

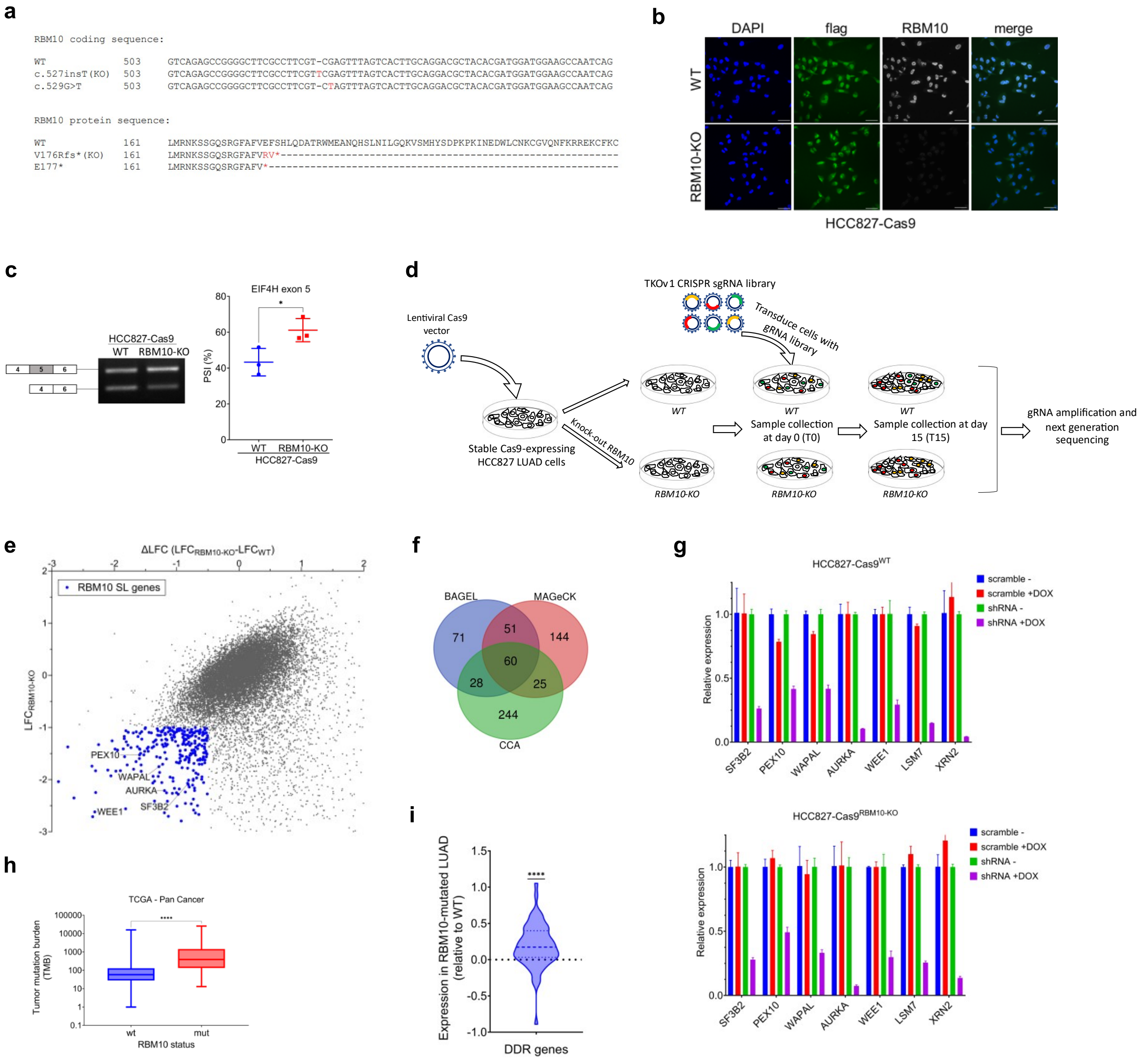


**Supplementary Figure 1: Functional validation of genome-wide CRISPR-Cas9 RBM10 SL screen. (a)** DNA and protein sequence alignment of the region in RBM10 exon 2 containing the position of the V176Rfs* mutation introduced using CRISPR-Cas9 system (highlighted in red) and the position of the E177* RBM10 cancer mutation. **(b)** Representative immunofluorescence microscopy images of HCC827-Cas9 isogenic cell lines co-stained for flag-Cas9 and RBM10. Nuclei are stained with DAPI. Scale bar, 50µm. **(c)** RT-PCR analysis of EIF4H exon 5 alternative splicing in HCC827-Cas9$^{WT}$ and HCC827-Cas9$^{RBM10\text{-}KO}$ cells. RNA was isolated from the indicated cell lines and analyzed by RT-PCR using primers flanking EIF4H exons 4-6. Left: Representative agarose gel image showing amplification of the two EIF4H variants that differ in exon 9 inclusion. Right: Percent-spliced-in (PSI) quantification of NUMB exon 9 inclusion. Data are presented as mean ± s.d. (n=3). Statistical significance was determined by unpaired t-test. **P<0.05*. **(d)** Schematic flowchart of the genome-wide CRISPR-Cas9 synthetic lethality screen conducted in isogenic RBM10-deficient and proficient HCC827 cells expressing flag-Cas9. **(e)** MAGeCK analysis results of CRISPR-Cas9 SL screen in WT and HCC827-Cas9$^{RBM10\text{-}KO}$ cells. Average gRNA log fold-change (LFC) for each gene in RBM10-KO is plotted against the LFC difference between RBM10-KO and WT cells. RBM10 SL genes are shown in blue. **(f)** Venn diagram showing the intersection of RBM10-SL genes identified using the 3 analysis methods: BAGEL, CCA, and MAGeCK. **(g)** Quantitative RT-qPCR analysis of TetON-shRNA conditional knockdown of the indicated genes in HCC827-Cas9$^{WT}$ (top) and HCC827-Cas9$^{RBM10\text{-}KO}$ (bottom). **(h)** Tumor mutation burden (TMB) analysis of The Cancer Genome Atlas (TCGA) database showing that tumors harboring RBM10 mutations display higher TMB levels. Statistical significance was determined by Mann-Whitney test. *****P<0.0001*. **(i)** Violin plot depicting the relative expression of DNA damage response signature genes in LUAD tumors harboring RBM10 mutations (n=38) compared to WT tumors with no RBM10 mutations (n=470). Data was retrieved from TCGA database. Thick line represents the median value and thin lines represent the quartiles. Statistical significance was determined by one-sample Wilcoxon test. *****P<0.0001*.

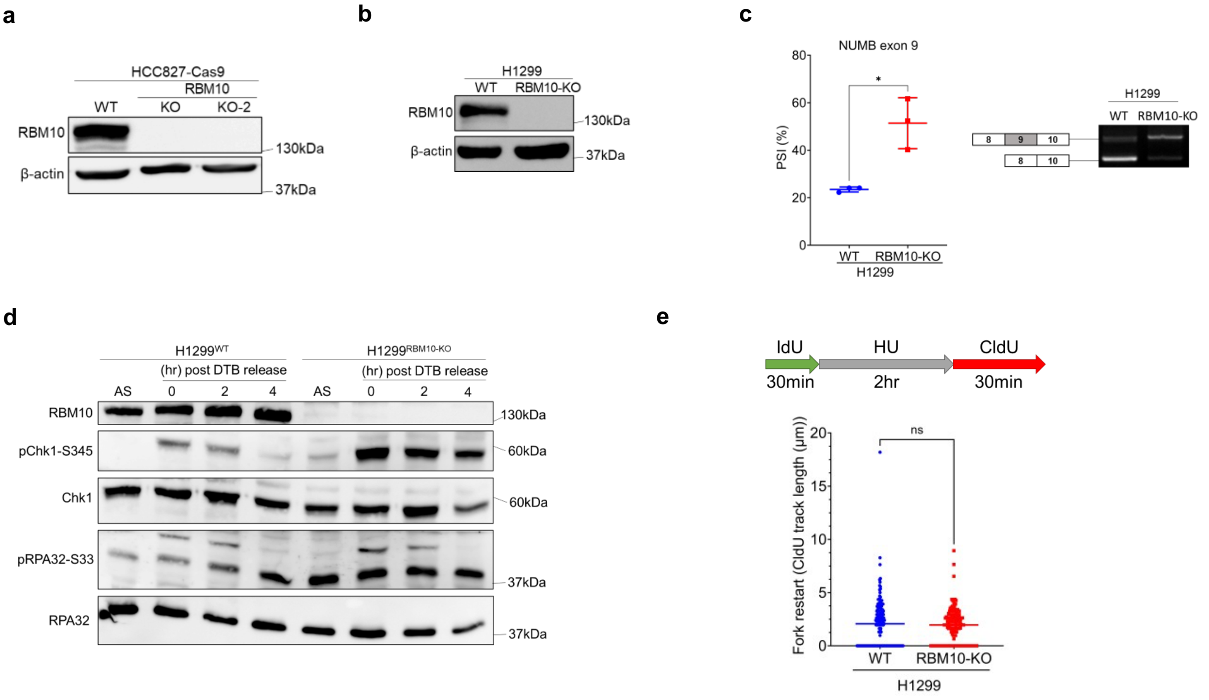


**Supplementary Figure 2: RBM10 promotes DNA replication fork progression and replication stress response. (a)** Immunoblot analysis for RBM10 protein expression in parental (WT) and HCC827-Cas9$^{RBM10\text{-}KO}$ cells. β-actin is used as a loading control. The positions of molecular weight markers are indicated to the right. **(b)** Immunoblot analysis for RBM10 protein expression in H1299$^{WT}$ and H1299$^{RBM10\text{-}KO}$ cells. β-actin is used as a loading control. The positions of molecular weight markers are indicated to the right. **(c)** RT-PCR analysis of NUMB exon 9 alternative splicing in WT and RBM10-KO H1299 cells. RNA was isolated from the indicated cell lines and analyzed by RT-PCR using primers flanking NUMB exons 8-10. Left: PSI (percent-spliced-in) quantification of NUMB exon 9 inclusion. Data are presented as mean ± s.d. (n=3). Statistical significance was determined by unpaired t-test. **P<0.05*. Right: Representative agarose gel image showing amplification of the two NUMB variants that differ in exon 9 inclusion. **(d)** H1299$^{WT}$ and H1299$^{RBM10\text{-}KO}$ cells were synchronized at S-phase using double-thymidine block (DTB) and subjected to immunoblot analysis at the indicated times after DTB release. AS=asynchronous cells. The positions of molecular weight markers are indicated to the right. **(e)** $H1299^{WT}$ and $H1299^{RBM10\text{-}KO}$ cells were analyzed by DNA combing following fork stalling by hydroxyurea (HU) as indicated above. The length of CldU+ region of IdU+ CldU+ replication tracks was measured (n = 150). Horizontal bars indicate the mean track length. Statistical significance was determined by Mann-Whitney test. ****$P<0.0001$.

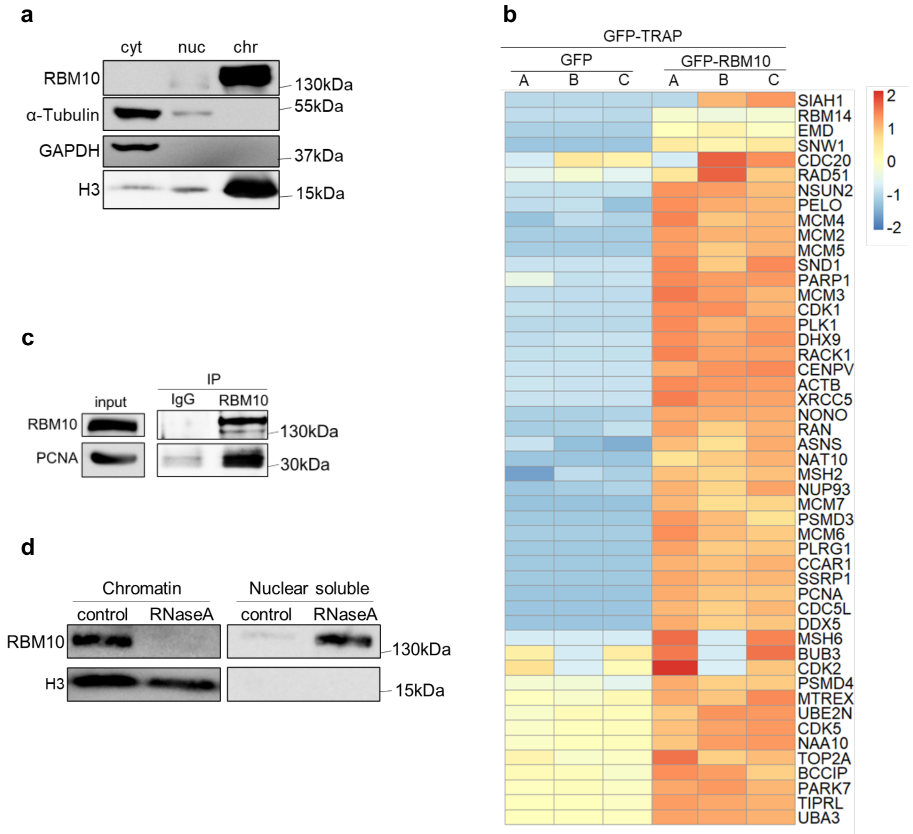


**Supplementary Figure 3: RBM10 interacts with replication fork components. (a)** Biochemical fractionation of HCC827 cells into cytosolic (cyt), nuclear-soluble (nuc), and chromatin-bound (chr) fractions followed by immunoblotting with the indicated antibodies. The positions of molecular weight markers are indicated to the right. **(b)** Heatmap summarizing RBM10-interacting proteins involved in cell cycle regulation, DNA damage response, and replication stress. Colors represent relative protein intensity of RBM10-interacting proteins in GFP-only and GFP-RBM10 samples across 3 replicates as identified by GFP-trap followed mass spectrometry. **(c)** Immunoprecipitation of endogenous RBM10 in H1299 cells. Whole-cell lysates from H1299 cells were subjected to immunoprecipitation using RBM10 antibody or IgG and subjected to immunoblot analysis with the indicated antibodies. The positions of molecular weight markers are indicated to the right. **(d)** Immunoblot analysis of RBM10 protein in chromatin-bound and nuclear-soluble fractions either treated with RNaseA or left untreated prior to extraction. The positions of molecular weight markers are indicated to the right.

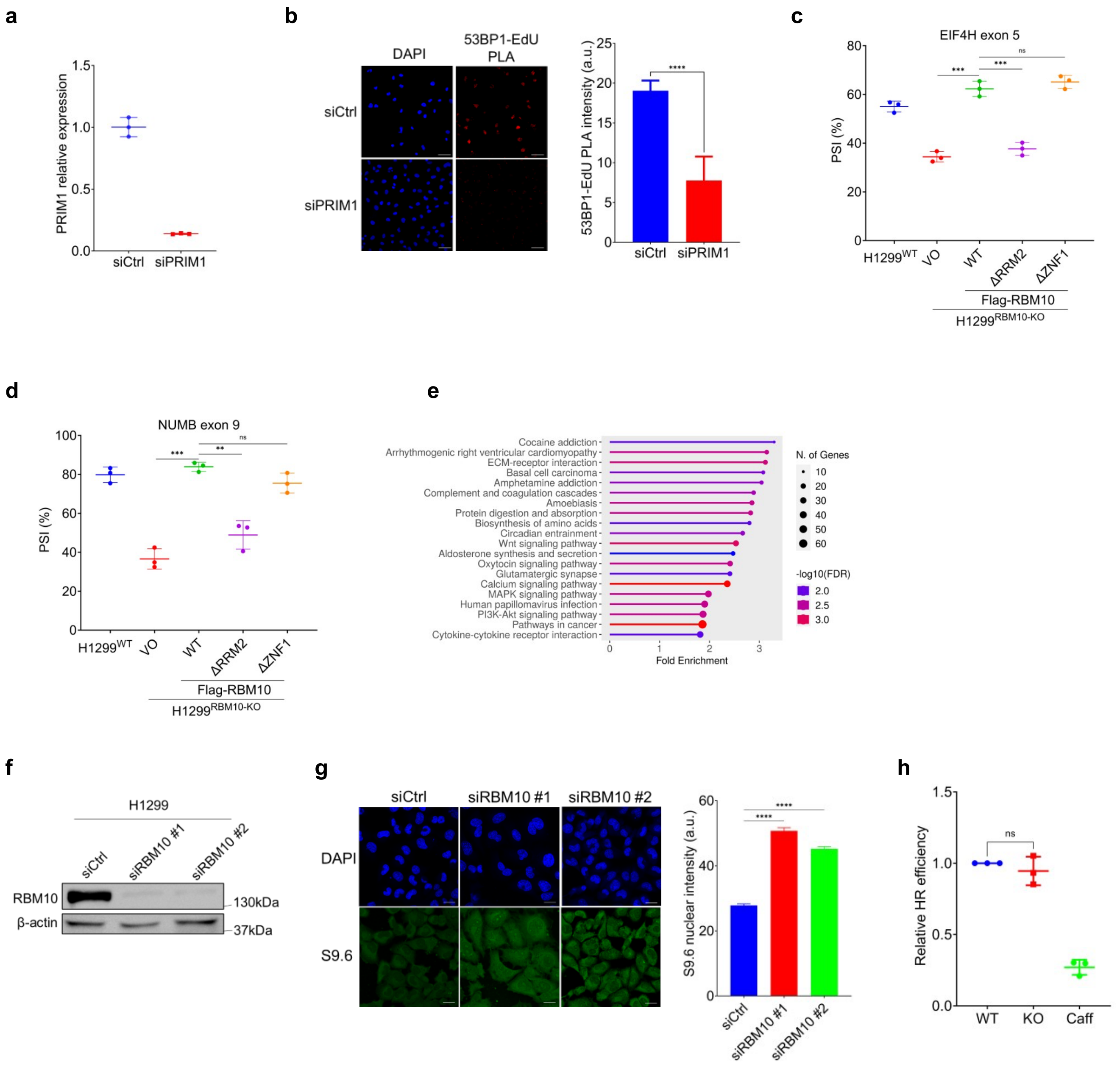


**Supplementary Figure 4: PRIM1-dependent RBM10 association with DNA replication forks promotes HDAC1 recruitment to limit replication stress. (a)** Quantitative RT-qPCR analysis of PRIM1 expression in H1299$^{WT}$ transfected with siRNA against PRIM1 (siPRIM1) or control siRNA (siCtrl) for 72 hr. **(b)** Left: Representative immunofluorescence microscopy images of 53BP1:EdU-biotin PLA in H1299$^{WT}$ cells transfected with siPRIM1 or siCtrl for 72 hr. DAPI is used to stain nuclei. Scale bar, 50μm. Right: Quantification of 53BP1-EdU-biotin PLA intensity per nucleus. Data are presented as mean PLA intensity per nucleus ± SEM (n>50 cells). Statistical significance was determined by Mann-Whitney test. *****P<0.0001*. **(c-d)** RT-PCR analysis of EIF4H exon 5 (c) and NUMB exon 9 alternative splicing in H1299$^{RBM10\text{-}KO}$ cells expressing Flag-RBM10$^{WT}$, Flag-RBM10$^{\Delta RRM2}$, Flag-RBM10$^{\Delta ZNF1}$, or vector only (VO). Data are presented as mean Percent-spliced-in (PSI) ± s.d. (n=3). Statistical significance was determined by unpaired t-test. ****P< 0.001;ns, not significant*. **(e)** Gene ontology analysis of differentially expressed genes identified by RNA-seq in three sets of cell lines sequenced in triplicates: parental and RBM10-deficient HCC827, parental and RBM10-deficient H1299, and HCC827 cells transfected with siRNA against RBM10 or control siRNA. Genes identified in at least 2 sets with absolute log-foldchange > 1 and p-value < 0.01 were used for gene ontology analysis. **(f)** Immunoblot analysis of RBM10 protein expression in H1299$^{WT}$ cells transfected with two siRNAs against RBM10 (siRBM10) or control siRNA (siCtrl) for 72 hr. β-actin is used as a loading control. The positions of molecular weight markers are indicated to the right. **(g)** Left: Representative immunofluorescence microscopy images of R-loops detected by S9.6 antibody in H1299$^{WT}$ cells transfected with two siRBM10 or siCtrl for 72 hr. DAPI is used to stain nuclei. Scale bar, 20μm. Bottom: Quantification of S9.6 signal intensity. Data are presented as mean intensity per nucleus ± SEM (n>100 cells) and representative of 3 independent experiments. Statistical significance was determined by Mann-Whitney test. *****P<0.0001*. **(h)** mClover assay to assess homologous recombination (HR) of endogenous DSBs at the LMNA gene. H1299$^{WT}$ and H1299$^{RBM10\text{-}KO}$ cells were transfected with plasmids expressing Cas9 endonuclease, LMNA-donor plasmid or Red-Monomer (MR) tag. 72 h after transfection, cells were harvested, and the percentage of GFP-positive cells from the total number of red cells was determined. Relative HR efficiency was determined by normalizing the percentage of GFP-positive cells to WT cells. Caffeine (Caff.) is used as a negative control. Data are presented as mean ± s.d. (n=3). *ns, not significant*.

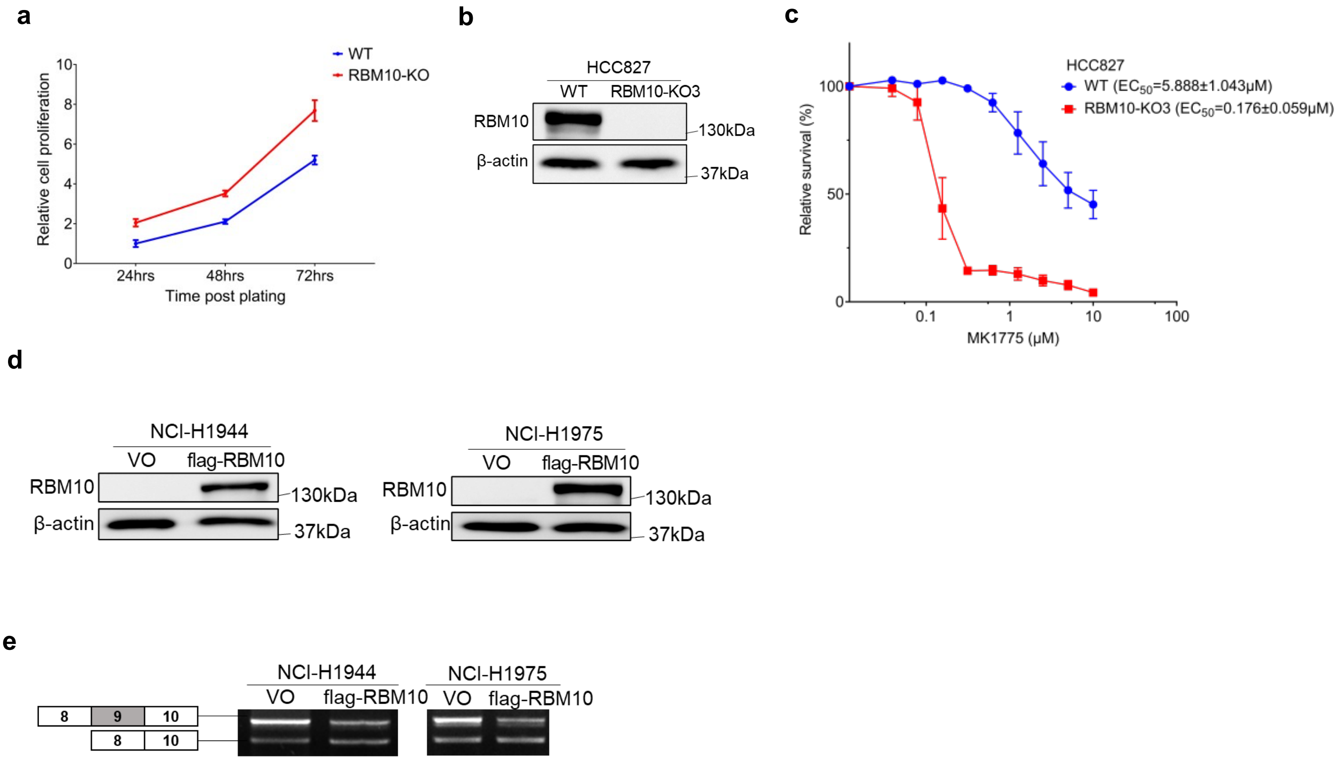


**Supplementary Figure 5: Characterization and functional validation of RBM10-deficient LUAD cells. (a)** Relative cell proliferation of parental (WT) and HCC827-Cas9$^{RBM10\text{-}KO}$ cells. Cell viability was measured at the indicated time post seeding and average cell viability (n=4) was normalized to the viability of WT cells. Data are presented as mean ± s.d. **(b)** Immunoblot analysis for RBM10 protein expression in naïve parental (WT) and HCC827$^{RBM10\text{-}KO}$ cells. β-actin is used as a loading control. The positions of molecular weight markers are indicated to the right. **(c)** Short-term cell viability assay and $EC_{50}$ determination in HCC827$^{WT}$ and HCC827$^{RBM10\text{-}KO}$ cells treated with increasing concentrations of MK1775. Data are presented as mean ± s.d. (n=3). **(d)** Immunoblot analysis for RBM10 protein expression in patient-derived RBM10-deficient NCI-H1944 (left) and NCI-H1975 (right) cells complemented with flag-RBM10-WT or empty vectors. β-actin is used as a loading control. The positions of molecular weight markers are indicated to the right. **(e)** RT-PCR analysis of NUMB exon 9 alternative splicing in NCI-H1944 and NCI-H1975 cells expressing flag-RBM10-WT or empty vectors. Representative agarose gel image showing amplification of the two NUMB variants that differ in exon 9 inclusion.

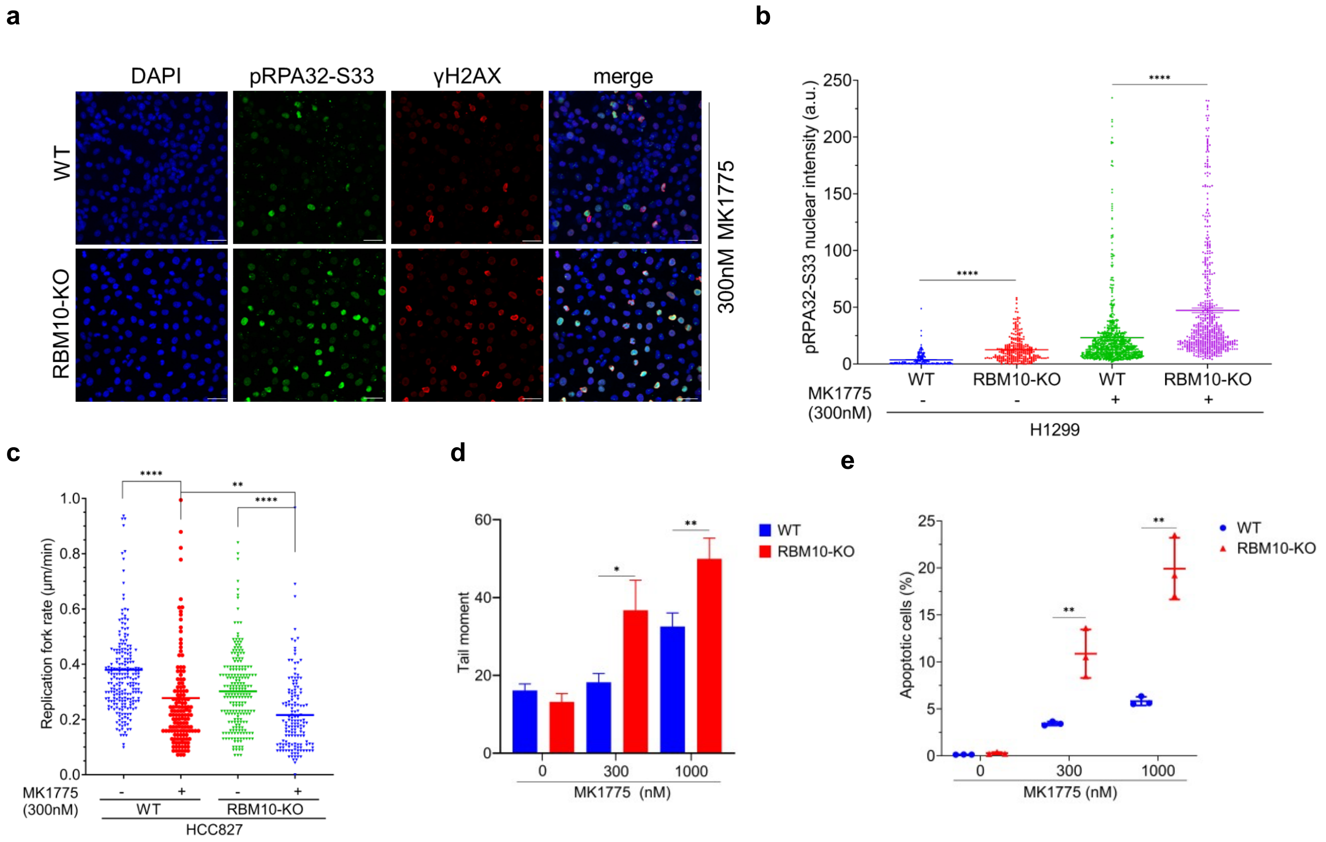


**Supplementary Figure 6: WEE1 inhibition leads to DNA damage accumulation in RBM10-deficient cells. (a-b)** Analysis of pRPA32-S33 intensity in H1299$^{WT}$ and H1299$^{RBM10\text{-}KO}$ cells treated with 300nM MK1775 for 24 hr. (a) Representative immunofluorescence microscopy image for pRPA32-S33 staining. DAPI was used to stain nuclei. Scale bar, 50µm. (b) Quantification of pRPA32-S33 staining. Data are presented as mean nuclear intensity ± SEM (n>200 cells for untreated cells, n>500 for MK1775-treated cells) and representative of 3 independent experiments. Statistical significance was determined by Mann-Whitney test. *****P<0.0001*. **(c)** DNA combing assay in HCC827-Cas9$^{RBM10\text{-}WT}$ and HCC827-Cas9$^{RBM10\text{-}KO}$ treated with MK1775. Replication fork speed (n>150) was measured in untreated cells and cells treated with 300nM MK1775 for 24 hr. Black bars represent mean value. Statistical significance was determined by Mann-Whitney test. ***P<0.01;****P<0.0001*. **(d)** Quantification of neutral comet assay in H1299$^{WT}$ and H1299$^{RBM10\text{-}KO}$ cells treated either with DMSO or the indicated concentrations of MK1775 for 24 hr. DNA damage is represented by comet tail moment. Data are presented as mean tail moment ± SEM (n>150) and representative of 3 independent experiments. Statistical significance was determined by Mann-Whitney test. **P<0.05;**P<0.01*. **(e)** Quantification of apoptosis in H1299$^{WT}$ and H1299$^{RBM10\text{-}KO}$ cells treated with the indicated concentrations of MK1775 for 24 hr. Cells were stained with Annexin V and propidium iodide and analyzed by FACS. Percentage of apoptotic cells was determined by %Annexin V–positive cells and data are presented as mean ± s.d. (n=3). Statistical significance was determined by unpaired t-test. ***P< 0.01*.

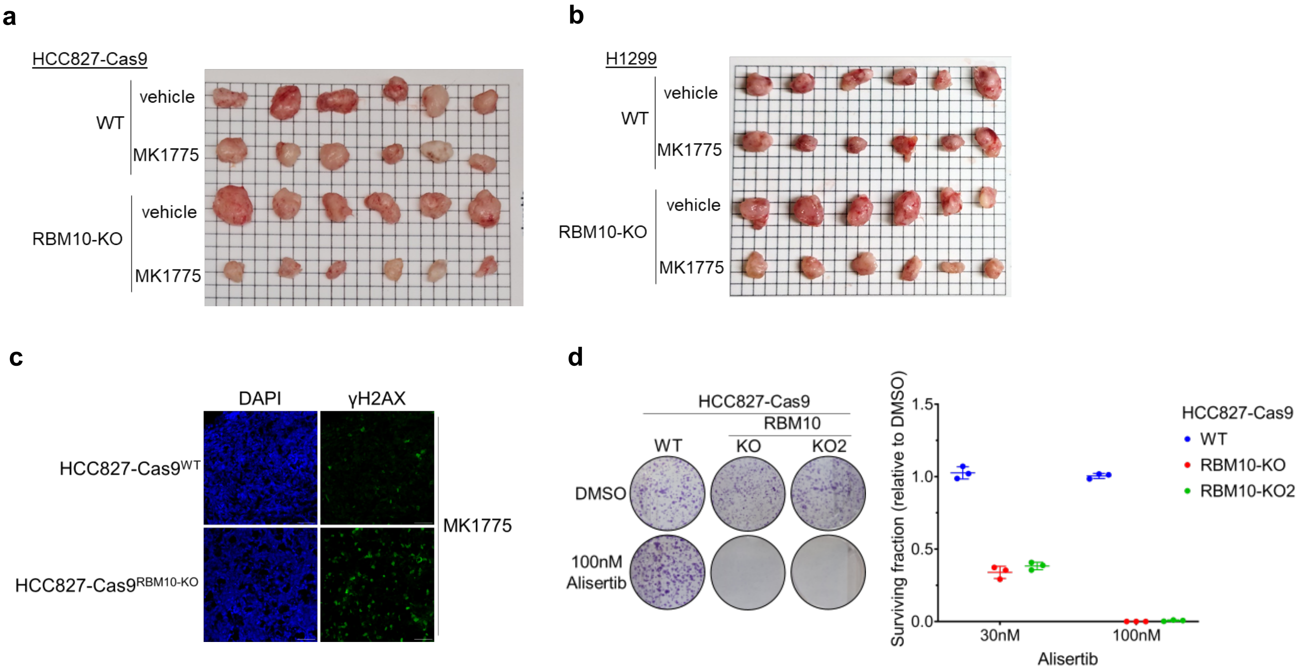


**Supplementary Figure 7: MK1775 inhibits the growth of RBM10-deficient xenograft tumors. (a-b)** Photographic images of parental (WT) and HCC827-Cas9$^{RBM10\text{-}KO}$ (a) and H1299$^{RBM10\text{-}KO}$ (b) xenografts treated with either MK1775 or vehicle. MK1775 was administered once daily at 40mg/kg for 15 days. Tumors from each group (n=6) were collected and imaged when the average tumor volume of vehicle-treated mice reached ~1500mm$^3$. Grid size, 0.5cm. **(c)** Representative immunofluorescence microscopy images of γH2AX staining in tumor sections from HCC827-Cas9$^{WT}$ and HCC827-Cas9$^{RBM10\text{-}KO}$ xenografts from MK1775-treated mice. DAPI is used to stain nuclei. Scale bar, 100μm. **(d)** Left, representative images of plates stained with crystal violet. Right, quantification of clonogenic survival. Data are presented as mean ± s.d. (n=3).

**Supplementary Tables:**

**Supplementary Table 1.** Excel file containing RBM10 CRISPR-Cas9 SL screen results in parental (WT) and RBM10-KO HCC827-Cas9. (a) gRNA read counts. (b) BAGEL analysis - gene essentiality scores are expressed as Bayes Factor (BF). (c) MAGeCK analysis results. (d) CCA analysis results.

**Supplementary Table 2.** Excel file containing mass spectrometry results of RBM10 GFP-trap. Protein identification and intensity quantification in 3 replicates of GFP-trap in HEK-293T cells expressing EGFP-only or EGFP-RBM10.

**Supplementary Table 3.** Excel file containing RBM10 differential expression results using DESeq2. (a) Sequencing read counts. (b) Parental (WT) and RBM10-KO H1299 cells. (c) WT and RBM10-KO HCC827 cells. (d) HCC827 cells transfected either with control siRNA or siRNA against RBM10.

**Supplementary Table 4. Primer sequences**

| gRNA and shRNA sequences | |
|---|---|
| Primer | Sequence |
| RBM10-gRNA1-F | CACCGCAAGTGACTAAACTCGACGA |
| RBM10-gRNA1-R | AAACTCGTCGAGTTTAGTCACTTGC |
| RBM10-gRNA2-F | CACCGTATCTCCCCGTATAGATCCG |
| RBM10-gRNA2-R | AAACCGGATCTATACGGGGAGATAC |
| Scramble-shRNA-F | CCGGGTGGACTCTTGAAAGTACTATCTCGAGATTTGACGGGTGGATAATCTGTTTTT |
| Scramble-shRNA-R | AATTAAAAACAGATTATCCACCCGTCAAATCTCGAGATAGTACTTTCAAGAGTCCAC |
| SF3B2-shRNA-F | CCGGGCTGATGTTGAGATTGAGTATCTCGAGATACTCAATCTCAACATCAGCTTTTTG |
| SF3B2-shRNA-R | AATTCAAAAAGCTGATGTTGAGATTGAGTATCTCGAGATACTCAATCTCAACATCAGC |
| PEX10-shRNA-F | CCGGCCACACTTGCAGGCTACCAGACTCGAGTCTGGTAGCCTGCAAGTGTGGTTTTTG |
| PEX10-shRNA-R | AATTCAAAAACCACACTTGCAGGCTACCAGACTCGAGTCTGGTAGCCTGCAAGTGTGG |
| AURKA-shRNA-F | CCGGCACATACCAAGAGACCTACAACTCGAGTTGTAGGTCTCTTGGTATGTGTTTTTG |
| AURKA-shRNA-R | AATTCAAAAACACATACCAAGAGACCTACAACTCGAGTTGTAGGTCTCTTGGTATGTG |
| WAPAL-shRNA-F | CCGGGCCCAATTTCAAACCAGATATCTCGAGATATCTGGTTTGAAATTGGGCTTTTTG |
| WAPAL-shRNA-R | AATTCAAAAAGCCCAATTTCAAACCAGATATCTCGAGATATCTGGTTTGAAATTGGGC |
| WEE1-shRNA-F | CCGGGCCAGTGATTATGAGCTTGAACTCGAGTTCAAGCTCATAATCACTGGCTTTTTG |
| WEE1-shRNA-R | AATTCAAAAAGCCAGTGATTATGAGCTTGAACTCGAGTTCAAGCTCATAATCACTGGC |
| LSM7-shRNA-F | CCGGCATCTTGGACTTGTCCAAGTACTCGAGTACTTGGACAAGTCCAAGATGTTTTTG |
| LSM7-shRNA-R | AATTCAAAAACATCTTGGACTTGTCCAAGTACTCGAGTACTTGGACAAGTCCAAGATG |
| XRN2-shRNA-F | CCGGTACATAGCTGATCGTTTAAATCTCGAGATTTAAACGATCAGCTATGTATTTTTG |
| XRN2-shRNA-R | AATTCAAAAATACATAGCTGATCGTTTAAATCTCGAGATTTAAACGATCAGCTATGTA |

| RT-PCR and qPCR primers | |
|---|---|
| Primer | Sequence |
| NUMB-Alt-F | GAAGTAGAAGGGGAGGCAGA |
| NUMB-Alt-R | GTCGGCCTCAGAGGGAGTA |
| EIF4H-Alt-F | TCACTTCGTGTGGACATTGC |
| EIF4H-Alt-R | CCTGCCCCCTAAGAAGTCAT |
| PEX10-RT-F214 | TTGAGCTGCTCTCAGATGTG |
| PEX10-RT-R303 | CTGGATGATGCTGACGTACT |
| SF3B2-RT-F2340 | GGCGATGGACGGAAGTGAGA |
| SF3B2-RT-R2464 | TCCGGCTCATAACCGTGGAC |
| WAPAL-RT-F3439 | CGGAATCGGCACTGTCTTGT |
| WAPAL-RT-R3584 | CGCTCTCGCTCAAGGAATAG |
| AURKA-RT-F1066 | TCCCACCTTCGGCATCCTAAT |
| AURKA-RT-R1250 | CAGTAAGACAGGGCATTTGCC |
| WEE1-RT-F1778 | AGAAGACCTTCAGCAATGGC |
| WEE1-RT-R1924 | GCCATCTGTGCTTTCTTGAG |
| LSM7-RT-F166 | CACTCCTCAACCTTGTGCTGGA |
| LSM7-RT-R296 | GCAGATTAGCACCACGGACGTG |
| XRN2-RT-F1568 | CAGTGAACCTGAGCCAGAGGAT |
| XRN2-RT-R1690 | GACTGCACAACTTTCCGACGGA |
| PRIM1-RT-F104 | TATCGCTGGCTCAACTACGGTG |
| PRIM1-RT-R219 | CACTCTGGTTGTTGAAGGATTGG |
| GAPDH-RT-F229 | CCAGGGCTGCTTTTAACTCT |

| | | |
|---|---|---|
| GAPDH-RT-R351 | GGTGCCATGGAATTTGCCAT | |
| **Cloning and mutagenesis primers** | | |
| Primer | Sequence | Usage |
| *Eco*RI-RBM10-F | GTCGGAATTCCATGGAGTATGAAAGACGTGGTGGTCGTG | Cloning RBM10 from pDest-RBM10-wt to pEGFP-C1 |
| *Bgl*II-RBM10-R | GGCAGATCTTCACTGGGCCTCGTTGAAGCGGGTCACCAT | |
| *Age*I-Flag-RBM10-F | GCACCGGTATGGATTACAAGGATGACGACGATAAGGAGTATGAAAGACGTGG | Subcloning RBM10 from pEGFP-C1-RBM10 to Lenti-Cas9-2A-Blast |
| *Bgl*II-RBM10-R-noStop | ATAAGATCTCTGGGCCTCGTTGAAGCGGG | |
| RBM10-dZnF1-F | AAGTCAGAGGCAGAGCAGAAGCTG | All-round PCR to generate pEGFP-C1-RBM10-ΔZNF1 |
| RBM10-dZnF1-R | ATTGATCTTGGGCTTGGGGTCACT | |
| RBM10-dRRM2-F | TTTGCCAAGGGTTCTAAGAGGGAC | All-round PCR to generate pEGFP-C1-RBM10-ΔRRM2 |
| RBM10-dRRM2-R | GTCATTGGCGTTCTCTGAGCTTGGC | |
| EcoRI-MCM5-F | GCATGGAATTCCATGTCGGGATTCGACGATCCTGGCAT | Used to clone MCM5 to generate p3X-Flag-MCM5 |
| BamHI-MCM5-R | GAAGGATCCTCACTTGAGGCGGTAGAGAACCTTGCG | |
